# Microbial genomes retrieved from High Arctic lake sediments encode for adaptation to cold and oligotrophic environments

**DOI:** 10.1101/724781

**Authors:** Matti O. Ruuskanen, Graham Colby, Kyra A. St.Pierre, Vincent L. St.Louis, Stéphane Aris-Brosou, Alexandre J. Poulain

## Abstract

The Arctic is currently warming at an unprecedented rate, which may affect environmental constraints on the freshwater microbial communities found there. Yet, our knowledge of the community structure and functional potential of High Arctic freshwater microbes remains poor, even though they play key roles in nutrient cycling and other ecosystem services. Here, using high-throughput metagenomic sequencing and genome assembly, we show that sediment microbial communities in the High Arctic’s largest lake by volume, Lake Hazen, are phylogenetically diverse, ranging from Proteobacteria, Verrucomicrobia, Planctomycetes, to members of the newly discovered Candidate Phyla Radiation (CPR) groups. These genomes displayed a high prevalence of pathways involved in lipid chemistry, and a low prevalence of nutrient uptake pathways, which might represent adaptations to the specific, cold (~3.5°C) and extremely oligotrophic conditions in Lake Hazen. Despite these potential adaptations, it is unclear how ongoing environmental changes will affect microbial communities, the makeup of their genomic idiosyncrasies, as well as the possible implications at higher trophic levels.

## Introduction

Climate change is transforming Arctic ecosystems: elevated temperatures and increasing precipitation have facilitated permafrost thaw (Romanovsky et al., 2010) and glacial melt (Lehnherr et al., 2018; Milner et al., 2017), leading to impacts on the functioning of aquatic and terrestrial ecosystems, and the natural services they provide. Microbes are major players in the biogeochemical cycling of organic matter and inorganic nutrients; as such studying contemporary Arctic microbial communities is critical for both documenting and predicting how these cycles might respond to ongoing and future environmental change. However, baseline microbial community data from the Arctic are still lacking because these environments are remote and challenging to study. Furthermore, in-depth study of microbial communities has only recently become possible with culture-independent methods like high-throughput sequencing (e.g., Shokralla et al., 2012). As Arctic environments are already responding to climate change at the watershed scale (Lehnherr et al., 2018), and these warming-related changes are projected to continue (IPCC, 2018), there is an urgent need to gather data on the current state of Arctic microbial communities and their function from which to compare in the future.

To date, culture-independent studies of microbial communities in Arctic lakes have been most often performed by amplifying and sequencing taxonomic markers like the 16S rRNA gene (e.g., Crump et al., 2012; Mohit et al., 2017; Ruuskanen et al., 2018; Stoeva et al., 2014; Wang et al., 2016, 2019). However, amplicon-based methods are subject to PCR amplification bias, which might alter the estimates of microbial community composition and diversity. Furthermore, while taxonomy based on a marker gene can be used to predict the functional potential of microbes (Louca et al., 2016), these predictions are purely hypothetical in the absence of functional data. Metagenomic sequencing enables the reconstruction of nearly complete Metagenome Assembled Genomes (MAGs) solely from environmental DNA (Zhou et al., 2015). Gene sequences coding for proteins derived from contiguous sequences in the metagenomes can also be used to reconstruct the functional potential of microbial communities in the sampled environment. For example, metagenomic sequencing enabled the discovery of the Candidate Phyla Radiation (CPR; Brown et al., 2015; Hug et al., 2016), consisting of uncultured, deeply branching lineages in bacteria, which had previously evaded detection in purely amplicon-based studies. After their initial discovery, CPR members have been found in a variety of environments, including the deep subsurface (Danczak et al., 2017), marine sediments (León‐Zayas et al., 2017), and hypersaline soda lakes (Vavourakis et al., 2018). Presence of CPR bacteria has also been reported in Arctic freshwater environments (Vigneron et al., 2019; Wurzbacher et al., 2017). While they appeared to be absent in 16S rRNA amplicon data, a metagenomic analysis of the same samples revealed them to be highly abundant (Vigneron et al., 2019). However, shotgun metagenomic data from lake sediment microbial communities in Arctic environments are still scarce (Wang et al., 2019), and both larger lakes and High Arctic lakes remain thus far uncharted by these methods. Such knowledge would definitely expand our understanding of Arctic and global microbial diversity.

To investigate this diversity of microbes and their metabolisms in understudied Arctic lakes, we analyzed shotgun metagenomes from sediment extracted DNA from Lake Hazen, the world’s largest High Arctic lake by volume. In a previous study using 16S rRNA gene amplicons, we hypothesized that taxonomically dissimilar sediment communities at different locations in the lake might be functionally similar (Ruuskanen et al., 2018). In the current study, we revisited this question by investigating the taxonomic diversity and functional potential of the sediment microbial community using metagenomics. We identified rarely studied organisms within the sediment, characterized the metabolic pathways that are over- and underrepresented in these reconstructed genomes (compared to reference data from online repositories as of June 2018), described the ecologically important nutrient cycles that are potentially present in the sediments, and identified which taxa might play key roles in them.

## Material and methods

### Sampling and chemical analyses

Lake Hazen (located within Quttinirpaaq National Park on northern Ellesmere Island, Nunavut, Canada; 81.8°N, 71.4°W) is 544 km^2^ in surface area, with a maximum depth of 267 m, making it the largest lake by volume north of the Arctic Circle. The area immediately surrounding the lake is a polar oasis with higher than average temperatures for similar latitudes. Temperatures over 0°C have been observed in this region on more than 80 days per year (Soper and Powell, 1985), which is likely due to thermal shielding by the Grand Land mountains in the northwestern portion of Lake Hazen’s watershed (France, 1993). The lake is primarily fed hydrologically by meltwaters of the outlet glaciers of the Grant Land Ice Cap, and has a single major outflow, the Ruggles River, which flows southeastwardly to the eastern coast of Ellesmere Island. Sediment cores were collected from two sites: Deep Hole [261 m] on the 4^th^ of August and Snowgoose Bay [49 m] on the 8^th^ of August 2016 (Figure S1). Sampling was conducted from a boat using an UWITEC (Mondsee, Austria) gravity corer with 86 mm inner diameter polyvinyl chloride core tubes. At both sites, triplicate cores were collected: one each for DNA extraction, porewater chemistry and microsensor measurements. While the extracted cores were up to 40 cm in length, the subsectioning was restricted to the topmost 6 cm in all cores, since microprobes cannot be pushed any deeper than this into the sediment cores. The core from which DNA were extracted was sectioned at 0.5 cm intervals in the field, preserved with LifeGuard™ (MO BIO, Carlsbad, CA), and stored at −18°C until DNA extraction. Because of logistical difficulties involved with sampling in the High Arctic, the sectioning equipment could only be cleaned with non-sterile lake water and bleach between each section before putting the complete sections into sterile 50 mL centrifuge tubes. Non-powdered nitrile gloves were worn while handling the samples. For porewater analyses (NH_4_^+^, NO_2_^−^/NO_3_^−^, SO_4_^2−^, TDP, Cl^−^), the core was similarly sectioned at 1 cm intervals and placed in sterile 50 mL centrifuge tubes. The sediment sections were centrifuged at 4500 rpm for 15 minutes to separate the sediments from the porewater. The supernatant was then filtered through 0.45-μm cellulose acetate syringe filters that were first rinsed with a bit of sample water. The remaining filtrate was stored in sterile 15 ml Corning polystyrene centrifuge tubes, and then immediately frozen at −18°C until analyses for NH_4_^+^, NO_2_^−^/NO_3_^−^, SO_4_^2−^, TDP and Cl^−^. Porewater chemical concentrations were determined using CALA-certified protocols at the Biogeochemical Analytical Service Laboratory (University of Alberta, Edmonton, AB, Canada). On the final core, 100-μm resolution microprofiles of O_2_, redox potential, and pH were measured using Unisense (Aarhus, Denmark) glass microsensors interfaced with the Unisense Field Multimeter immediately upon return to camp. Cores were maintained at ambient temperatures (~4°C) throughout profiling. The microprofiles of these cores have also been described in an earlier study (St. Pierre et al., 2019).

### DNA extraction and sequencing

For the extraction of environmental DNA, triplicate ~0.5 g (wet weight) subsamples were taken from three intervals per sediment core (Deep Hole: 0.0 – 0.5 cm, 1.0 – 1.5 cm, 2.0 – 2.5 cm; Snowgoose Bay: 0.0 – 0.5 cm, 1.5 – 2 cm, 3.0 – 3.5 cm). The samples were first washed once with a saline buffer (10 mM EDTA, 50 mM Tris-HCl, 50 mM Na_2_HPO_4_·7H2O at pH 8.0) to remove inhibitors (Poulain et al., 2015; Zhou et al., 1996), and then DNA was extracted from the samples (and a negative reagent control) with the DNeasy PowerSoil Kit (Qiagen, Hilden, Germany) according to the manufacturer’s instructions. Sample manipulations and extractions were conducted with sterilized equipment in a laminar flow cabinet (HEPA 100). The quality of the DNA was checked with a NanoDrop 2000 (Thermo Fisher Scientific, Wilmington, DE, USA) and through confirming the amplification of the *glnA* gene from the extracts (with *E. coli* DNA as the positive and sterile H_2_O as the negative control) by PCR and gel electrophoresis (see SI text). The *glnA* gene was chosen as the control for DNA quality because its genomic copy number should be lower than that of the 16S rRNA gene (Stoeva et al., 2014) and a positive result would better confirm the quality of the DNA. The triplicate DNA extracts were then combined for each core horizon. Library preparation and sequencing was completed by Genome Quebec (Montreal, QC, Canada) with Illumina HiSeq 2500 PE125 in triplicate lanes for each sample.

### Preprocessing high-throughput sequencing data

The forward and reverse reads were trimmed and filtered for size and quality with Trimmomatic 0.36 (Bolger et al., 2014) and the data from all six samples were co-assembled with Megahit v1.1.2 (Li et al., 2015). Anvi’o v4 (Eren et al., 2015) was used for database management, following their standard metagenomic workflow. Briefly, reads were mapped to the contigs with Bowtie 2 (Langmead and Salzberg, 2012) and contigs longer than 1 kbp were binned with CONCOCT 0.4.1 (Alneberg et al., 2014). Open reading frames were identified with Prodigal (Hyatt et al., 2010), and functional annotations were inferred based on six reference systems: NCBI’s Cluster of Orthologous Genes (COG; Galperin et al., 2015; Tatusov et al., 2003) was run within Anvi’o using DIAMOND (Buchfink et al., 2015) for the protein alignments using the default E-value cutoff of 0.001. Following this, Pfam (Finn et al., 2014), TIGRFAM (Selengut et al., 2007) and Gene Ontology (GO; Ashburner et al., 2000; The Gene Ontology Consortium, 2017) annotations for proteins were added with InterProScan 5.29-68.0 (Jones et al., 2014). Briefly, for both the Pfam and TIGRFAM reference systems, the query protein sequences were searched with HMMER3 (Eddy, 2009) against the respective hidden Markov model databases. Hits for individual query proteins were filtered based on curated model-specific cut-offs and, in the case of Pfam, lower-scoring hits in the same Pfam clan were removed (Jones et al., 2014). Finally, GO terms were associated with the proteins within InterProScan through cross-referencing the Pfam and TIGRFAM annotations with the InterPro database (Hunter et al., 2012).

These bins that were assembled automatically from the contigs were then manually refined in Anvi’o to < 10% contamination (also named “redundancy” in the Anvi’o documentation), based on single copy genes following Campbell et al. (2013) for bacteria, and Rinke et al. (2013) for archaea. These contamination estimates were cross-compared against the lineage-specific marker genes from CheckM v1.0.11 (Parks et al., 2015), and bins that still had CheckM contamination > 10% were further refined manually by splitting. Final completion values for the refined genome bins were also estimated with CheckM. Read coverages of the MAGs in each sample were calculated with Anvi’o, normalized to total number of reads in each sample, and finally scaled to binned reads in each sample. Genome bins were analyzed for the presence of KEGG (Kanehisa et al., 2017) and MetaCyc (Caspi et al., 2018) pathways by mapping GO terms of each bin to Enzyme Classification (EC) categories and reconstructing parsimonious pathways with MinPath (Ye and Doak, 2009). The reconstructed genomes were also analyzed for the presence of marker genes (Table S1) and MetaCyc pathways (Table S2) for core elemental cycles (C, N, P, and S), together with metal regulation and homeostasis genes. Abundances of individual MetaCyc pathways were calculated by summing the sample-wise normalized abundances of each MAG indicated to contain the pathway. The abundance of the individual marker genes in the separate samples was calculated from gene-level read coverages in Anvi’o, followed by normalization to total reads in a sample.

To estimate the robustness of the MAG-based community composition in the samples, reads identified as 16S rRNA genes were extracted from the dataset, and assembled with MATAM (Pericard et al., 2018). The 16S rRNA gene contigs were then classified against the SILVA 128 NR95 database (Quast et al., 2013) with the RDP Classifier (Wang et al., 2007). The abundances of the operational taxonomic units (individual 16S rRNA contigs) in MATAM were calculated as the proportion of 16S rRNA reads mapping to each contig per sample.

For phylogenetic and taxonomic comparisons of MAGs to reference data, all non-redundant (one per species) complete bacterial and archaeal genomes, and all the available Candidate Phyla Radiation (CPR; Brown et al., 2015; Hug et al., 2016) genomes, were downloaded from the NCBI GenBank database (Benson et al., 2012). As of June 2018, this comprised 3,362 bacterial, 240 archaeal and 3,561 genomes for CPR. A further 71 CPR genomes were added from Danczak et al. (2017). Open reading frames were used if available, or they were identified de novo with Prodigal. Functional annotations for genes and pathways in these genomes were performed de novo as described above for MAGs. For assessing the phylogeny, sequences of 16 ribosomal proteins were extracted from both the NCBI genomes and our MAGs that were > 70% complete (following Hug et al., 2016; Table S3). The sequences were aligned per ribosomal protein with MAFFT 7.402 (Katoh and Standley, 2013) using translated protein sequences, and back-translated to the original nucleotide sequences with TranslatorX (Abascal et al., 2010). Badly aligned sequences were removed, and the alignments were trimmed with trimAl 1.2rev59 using the ‘-gappyout’ mode (Capella-Gutiérrez et al., 2009). Phylogenetic trees were constructed from each ribosomal protein with FastTree 2.1.9, compiled with double precision to estimate accurately short branch lengths (as recommended in the manual), and using the GTR + Γ model of sequence evolution (e.g., Aris-Brosou and Rodrigue, 2012). Sequences with unexpectedly long branches in the individual ribosomal protein trees were then removed with treeshrink (Mai and Mirarab, 2018) with tolerance of false positives ‘--quantiles’ set to 0.01. The trimmed alignments were concatenated for each genome with gap characters added for missing ribosomal proteins. NCBI genome entries with more than 25% gaps and MAGs with more than 50% gaps over the full alignment were then removed.

The higher cut-off on gap proportion for MAGs (50%) compared to the reference genomes (25%) was used to enable inclusion of a higher number of MAGs in the downstream analyses. However, together with the low cut-off used in the binning step (all >1 kbp contigs included), the phylogenetic uncertainty of our taxonomic assignments could have been increased. Thus, we calculated the pairwise phylogenetic distance of each MAG against the reference genomes for each of the ribosomal protein trees with the ‘cophenetic’ function from ape (Paradis et al., 2004), and the number of incidences of each reference genome in the ten closest genomes for each MAG. The correlation between the proportional incidence of the most commonly found reference genome in each MAG was then compared against the number of different contigs containing ribosomal proteins in them with a least-square linear regression, where the number of contigs was log10-transformed. Finally, the phylogenetic tree containing both the MAGs and the reference genomes was constructed with FastTree 2.1.9 under the GTR + Γ model of sequence evolution, as above.

### Taxonomy analysis and functional potential of the microbial community

For further data analysis, only MAGs with < 50% gaps over the complete alignment were preserved (*n* = 55; Table S4). The phylogenetic tree containing the MAGs and reference genomes was visualized with ggtree (Yu et al., 2017), and annotated based on NCBI taxonomy. The taxonomy of the MAGs was manually annotated based on the lowest level in a monophyletic clade starting from a node with a support value of at least 0.5 (full tree is included in the SI both as a PDF and a Newick tree file). For MAGs with uncertain taxonomy assignments at the phylum level, 16S rRNA sequences were extracted from their genomes (> 200 bp; *n* = 2) with ssu-align 0.0.1 (Nawrocki and Eddy, 2010). Also, the 16S rRNA sequences of their closest relative reference genomes based on the ribosomal protein tree were extracted. The MAG 16S rRNA sequences were matched against the NCBI’s nr database with BLASTN 2.8.1+ (Zhang et al., 2000) and the top 50 hits for each query were downloaded. These collections consisting of 16S sequences from the MAG itself, *Thermanaerovibrio acidaminovorans* DSM 6589, Candidatus *Caldatribacterium saccharofermentans* OP9-77CS, *Acetomicrobium hydrogeniformans* (NCBI Reference Sequence: NR_116842.1), and the 50 top matches from the BLASTn searches were then aligned and trees built with the Silva Alignment and Tree service (Pruesse et al., 2012), using FastTree under the GTR + Γ model of sequence evolution. The trees were then rooted using the *Acetomicrobium hydrogeniformans* 16S rRNA gene as the outgroup.

Phyloseq 1.24.0 (McMurdie and Holmes, 2013) was used to manage abundance and phylogenetic data. Phylum-level microbial community composition was qualitatively compared between the 16S rRNA assembly and the genome assembly of 55 MAGs, where taxonomy assignments were based on the tree of ribosomal proteins. Differences in the phylum-level 16S rRNA based taxonomy between the samples were also assessed by fitting a generalized linear model with a quasi-binomial link function (using ‘glm’; R Core Team, 2018) to the data. To examine patterns in community structure and function of the MAGs, between-sample distances were first calculated for genome phylogeny using patristic distances with Double Principal Coordinate Analysis (DPCoA; Pavoine et al., 2004). For 16S-based taxonomy, and functional pathways, Bray-Curtis dissimilarities were calculated in vegan 2.5.2 (Oksanen et al., 2018). The distance matrices were ordinated with NMDS, and envfit (Oksanen et al., 2018) was used to correlate the ordinations with sample physicochemistry. To further compare clustering patterns, the distance matrices were dimensionally reduced with *t*-Distributed Stochastic Neighbor Embedding (*t*-SNE; van der Maaten and Hinton, 2008) with the R package rtsne 0.13 (Krijthe and van der Maaten, 2017) and ‘perplexity’ set to 1.6. Clusters were then detected with Hierarchical Density-Based Spatial Clustering of Applications with Noise (HDBSCAN; Campello et al., 2013) in dbscan 1.1.2 (Hahsler et al., 2018) with ‘minPts’ set to 2.

The marker genes and MetaCyc pathways were classified to environmentally relevant cellular processes (Carbon (C), Nitrogen (N), Sulfur (S), Phosphorus (P), and toxic metal cycling) and these were divided into categories (Table S5). Some pathways were deemed to be misclassified based on their usual taxonomic range specified in the MetaCyc pathway descriptions and subsequently removed from the data. The proportion of genomes that had at least one single marker gene or pathway for a process were quantitatively compared between the MAGs (*n* = 55) and reference genomes subset to phyla shared with the MAGs (*n* = 2,486). The homogeneity of pathway abundances between MAGs to the reference genomes was assessed with a Pearson’s χ^2^ test where *P*-values were estimated based on 10,000 permutations. Differentially abundant processes for each comparison, which contributed more than their equal share (out of *n* = 46 processes) to the total *X*^2^ score, were visualized with heatmaps of their Pearson residuals. Finally, the gene-level read coverages (normalized to total number of reads per sample) of N, S, P, and toxic metal processing marker genes in the Lake Hazen sediments were compared between the two sites and between oxic (> 0.0 mg L^−1^) and anoxic (0.0 mg L^−1^) samples. The significances of the differences were assessed with pairwise *t*-tests, using ‘mt’ (False Discovery Rate - based correction for multiple testing) from Phyloseq 1.24.0 (McMurdie and Holmes, 2013). To identify the most abundant MAGs and phyla for each process across the samples, the read coverage of each of the 55 MAGs (relative to total number of reads per sample) was averaged over all six samples (Table S6). The relative abundances of the MAGs were also summed together at the phylum level (or class for Proteobacteria) by processes identified through MetaCyc pathways or individual marker genes (Table S7). The most abundant phylum in each process was then identified for inclusion in the flow-charts of the N, S, and Hg cycles (Figure 5).

Primary sequence data produced in this study is available in the NCBI Sequence Read Archive and the 55 medium-quality MAGs are available in GenBank (https://www.ncbi.nlm.nih.gov/bioproject/PRJNA525692). The geochemical data are available in the NSF Arctic Data Center online repository (https://arcticdata.io/catalog/#view/doi:10.18739/A2SJ19Q9P). Shell scripts, and code used in R 3.5.2 (R Core Team, 2018), are available in the supplementary material and through GitHub (https://github.com/Begia/Hazen-metagenome).

## Results and discussion

### Reconstruction of Metagenome Assembled Genomes (MAGs)

A total of 115.6 Gbp of sequence in 685 million reads were obtained from the six samples (Figure S2). The reads were co-assembled into 5.3 million contigs consisting of 3.7 Gbp of non-redundant sequences. The longest contig was 408.1 kbp and the N50 was 748 bp. The performance of our assembly was slightly better than when the same assembler was used for a complex soil metagenome (Howe et al., 2014; Li et al., 2015), but comparable to other studies of sediments with the same assembler (e.g., Carr et al., 2018; Lau et al., 2018). We sorted the contigs into 1488 bins with < 10% contamination, of which 146 were at least ‘medium-quality draft’ assemblies with > 50% completion (after Bowers et al., 2017). The remaining bins were ‘low-quality draft’ assemblies at < 50% completion (*n* = 1342). On average, 58% of all reads per sample were included in the 1488 bins.

Of these 146 medium-quality draft genomes, 68 were found to be > 70% complete, of which only a subset of 55 MAGs had > 50% of the nucleotides in the concatenated alignment of ribosomal proteins, deemed to be the minimum amount of information for placing them in the genome tree. On average, these 55 MAGs had a completeness of 88.2% based on single copy genes, a G/C ratio of 52.9, an N50 of 22 kbp, a genome size of 3.1 Mb, and contained 3,117 genes with a coding density of 90% (Table S4). Note that we used a 1 kbp lower bound on contig size during the binning process, essentially to increase the number of contigs in this step, which is somewhat lower than has recently been used (e.g., 2.5 kbp and 5 kbp in Delmont et al., 2018). However, only eight MAGs (14.5% of the total) appeared to have low N50 values (2.3 - 5 kbp), and the phylogenetic uncertainty in the taxonomic assignments of the MAGs was not correlated to the number of contigs containing ribosomal proteins in each MAG (*P* = 0.9). Thus, while this lower 1 kbp cut-off that we used in binning and taxonomy assignment of the MAGs might have increased their fragmentation, this liberal cut-off did not increase the uncertainty of their phylogenetic placement. These are key considerations for the reliability of the downstream analyses, because both the reconstructions of the metabolic pathways of the 55 MAGs (with MinPath) and their phylogenetic placement (with ribosomal proteins) are based on the collection of annotated genes contained in the individual MAGs.

### Differential recovery of MAGs compared to 16S rRNA gene data

The most common phyla (or class for proteobacteria) in the MAGs were Verrucomicrobia (14%), Betaproteobacteria (13%), Alphaproteobacteria (12%), Planctomycetes (10%) and Actinobacteria (9%; Figure 1a). To quantify the reliability of this taxonomic profile, we also binned the raw reads separately with the MATAM pipeline specifically designed to identify 16S rRNA genes (Pericard et al., 2018). This analysis, performed with the default settings of the pipeline, produced 166 OTUs which covered at least 500 bp of the complete 16S rRNA gene, about a third of its length. Among these OTUs, the most common bacterial and archaeal phyla were Betaproteobacteria (20%), Woesearchaeota (13%), Alphaproteobacteria (10%), Actinobacteria (8%) and Deltaproteobacteria (5%; Figure 1b). As a result, the 55 medium-quality MAGs represented a differential recovery of the total (16S rRNA reads-derived) microbial community, apparently missing at least two major taxonomic groups, Archaea and Firmicutes. This may be due to challenges inherent to identifying taxa based on 16S data and discrepancies between the NCBI and the SILVA databases (Parks et al., 2018), as well as at least two additional issues

**Figure 1.**
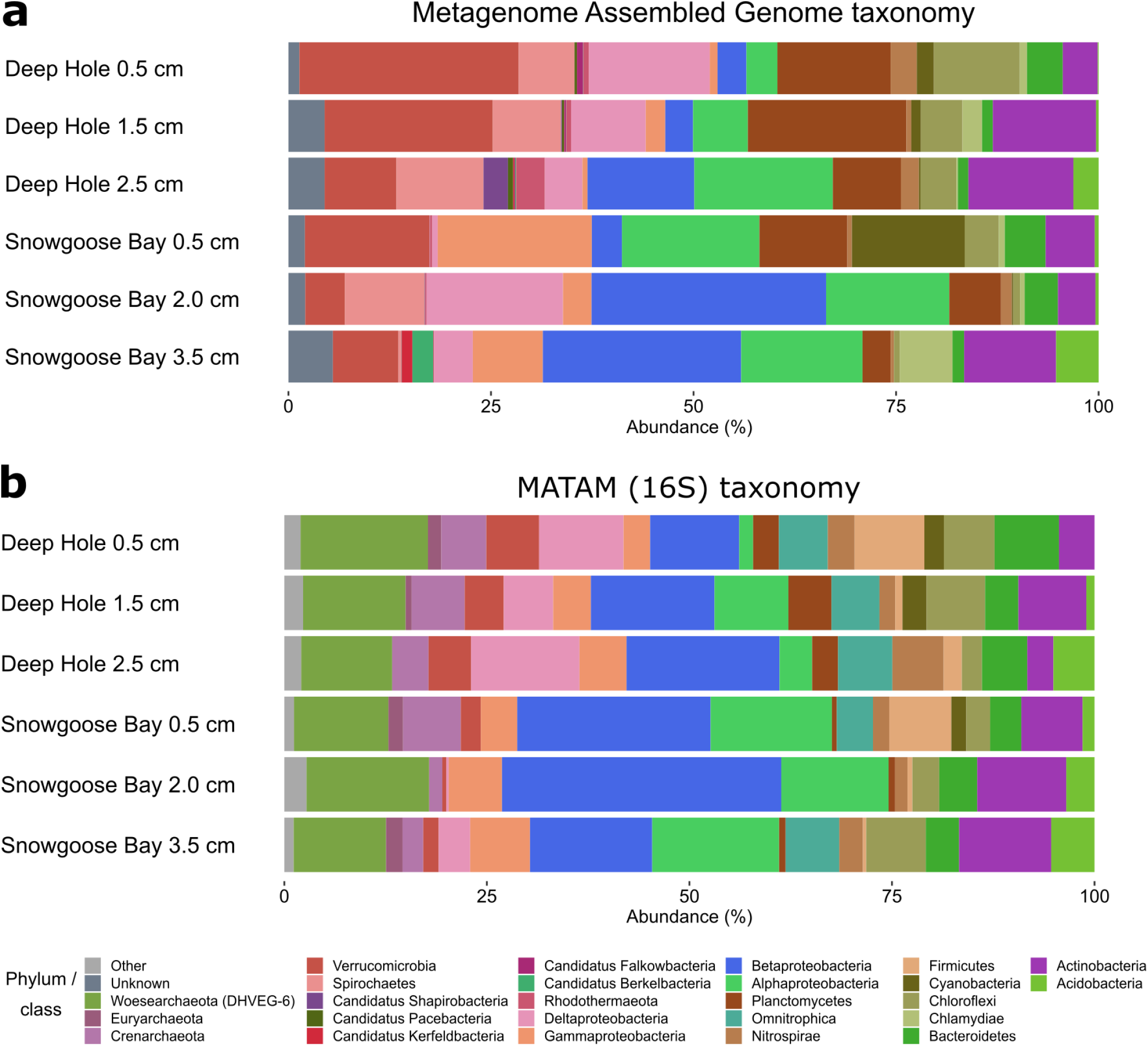
Composition of the microbial communities in the samples from Lake Hazen sediments. (**a**) Medium-quality draft MAGs (n = 55) with manual taxonomy assignments based on reference genomes in a phylogenetic tree constructed from ribosomal proteins. The phylum Proteobacteria is here subdivided into classes. (**b**) Raw reads binned into 16S rRNA contigs (n = 166) with MATAM and automatic taxonomy assignments from the SILVA 128 NR95 database.

First, while we identified 25 archaeal bins in the assembly, only six of them were over 10% complete and none were more than 70% complete. It should be noted that the six sequenced samples had similar community compositions based on the 16S rRNA reads-derived data (Figure 1b; all *P* = 1.0 at the phylum-level), which might have increased the binning difficulty because it utilizes differences in read coverages of contigs between samples (e.g., Alneberg et al., 2014). Second, Firmicutes appeared to be underrepresented among the assembled bins. Only nine low-quality Firmicutes genomes were assembled, of which seven were between 10% and 50% complete, and the rest below 10% complete. This underrepresentation of Firmicutes in the assembled genomes might have been caused by their endosporulation affecting DNA extraction, making them detectable only by marker genes (Filippidou et al., 2015). Despite the difficulties related to the metagenomic assembly, the taxonomic composition of the microbial community in Lake Hazen sediments was very similar to our previous study of the same sites in spring 2015 (Ruuskanen et al., 2018). However, several groups that were found here to be highly prevalent (Archaea, Omnitrophica) were likely missed in the earlier study because of PCR selection bias (Suzuki and Giovannoni, 1996). The community was also similar to previous metagenomic studies of oligotrophic lake sediments (e.g., Wang et al., 2016). One notable exception to previous studies (e.g., Rautio et al., 2011) was that Cyanobacteria were much less common in Lake Hazen. This is likely due to most other arctic studies having focused on shallow thaw ponds, whereas Lake Hazen is ~260 m deep, with light penetration restricted to the upper ~25 m of the water column. Furthermore, other physicochemical characteristics of Lake Hazen (such as ultra-oligotrophy) may not be favorable to Cyanobacteria (St. Pierre et al., 2019), which might affect their abundance in sediments as well. It should also be noted, that at least the phylum-level microbial community composition in Lake Hazen sediments appeared to be stable from 2015 spring to 2016 summer, despite high sedimentation rates in summer of 2015 resulting from enhanced glacial melt (St. Pierre et al., 2019).

To understand how the physicochemistry of the lake sediments (Figure S3) can drive community structure and function, we used ordination and clustering methods to compare the samples (for an in-depth analysis of Lake Hazen contemporary limnology, see St. Pierre et al., 2019), first based on the 16S-derived data. Here, the sites from Lake Hazen did not have significantly different microbial communities (based on 10,000 permutations; *P* = 0.10), and Cl^−^ concentration was the only chemical variable with a significant linear correlation with the differences in microbial community structure (*P* = 0.03; Figure S4). However, the Cl^−^ concentration was very low, varying within a narrow range (0.11 – 0.27 mgL^−1^), and also covarying with both pH and SO_4_^2−^ concentration, which are both known to influence the community structure in Lake Hazen (Ruuskanen et al., 2018). Furthermore, we observed no significant differences in community structure along the redox gradient (*P* = 0.77) or O_2_ concentration (*P* = 0.23) – which is partly consistent with a previous study of Lake Hazen sediments that showed that the microbial community was not constrained by oxygen concentration, but that the redox gradient was associated with community structure (Ruuskanen et al., 2018). This discrepancy is however not unexpected, as the range of redox potentials of the samples in the current study (Figure S3) was much narrower than in the previous study. Furthermore, the O2 concentration likely plays some role in the community structure and function in Lake Hazen sediments, but the gradient can be extremely steep (Figure S3; Ruuskanen et al., 2018), and differences could be only seen with comparisons of much thinner sediment horizons-.

To assess the validity of these 16S-derived results, we turned to the 55 MAGs, which also showed no significant differences in terms of community structure among samples, or in terms of correlations between community structure and physicochemistry (all *P* > 0.10). Similarly, a clustering analysis was unable to fully separate the sites based on these 55 MAGs (Figure S5A), but not on the 16S rRNA contig data that exhibited more differences among sites (with identical settings as for the MAGs; Figure S5B); note that these clustering analyses are by nature however qualitative, as no statistical tests were performed. Finally, we saw no separation of the samples in clustering when using functional pathway data from the 55 MAGs (derived with MetaCyc; Figure S6). This was also likely due to the small differences in the community composition of the two sites. As the communities appeared to be similar at both sampling sites, we pooled the data from each site together for all downstream analyses, in order to assess the extent of phylogenetic and potential metabolic microbial diversity in those lake sediments.

### Lake Hazen sediments harbor phylogenetically diverse bacteria

To place the 55 MAGs in a phylogenetic and taxonomic context, we reconstructed a tree adding 5,942 reference genomes to our MAG collection (Figure 2). The analysis showed that five of the MAGs (9%) likely represented CPR bacteria from previously known, NCBI-classified, candidate phyla (namely: Kerfeldbacteria, Falkowbacteria, Berkelbacteria, Pacebacteria and Shapirobacteria). This result highlights the importance of sequencing samples from rarely studied environments, such as High Arctic lakes. Two of these MAGs displayed similar characteristics to previously studied CPR (Hug et al., 2016), such as small genome size (< 1.3 Mbp) and a low number of genes (1309 and 1255 open reading frames; Table S4). In addition, three MAGs could not be classified to any known phylum by either their ribosomal proteins, or their partial 16S rRNA gene sequences (SH-like aLRT support < 50%). We note however that among these three unclassified MAGs, LH_MA_65_9 was likely related to uncultured bacteria close to candidate division OP11 (Figure S7), and LH_MA_57_9 was likely related to bacteria close to candidate division OP10 (Figure S8), a division mostly sampled from lake sediments. For example, the sediments of Upper Mystic Lake (Massachusetts, USA, www.ncbi.nlm.nih.gov/nuccore/DQ166697.1) contained the closest match to LH_MA_65_9 (99% BLAST identity).

**Figure 2.**
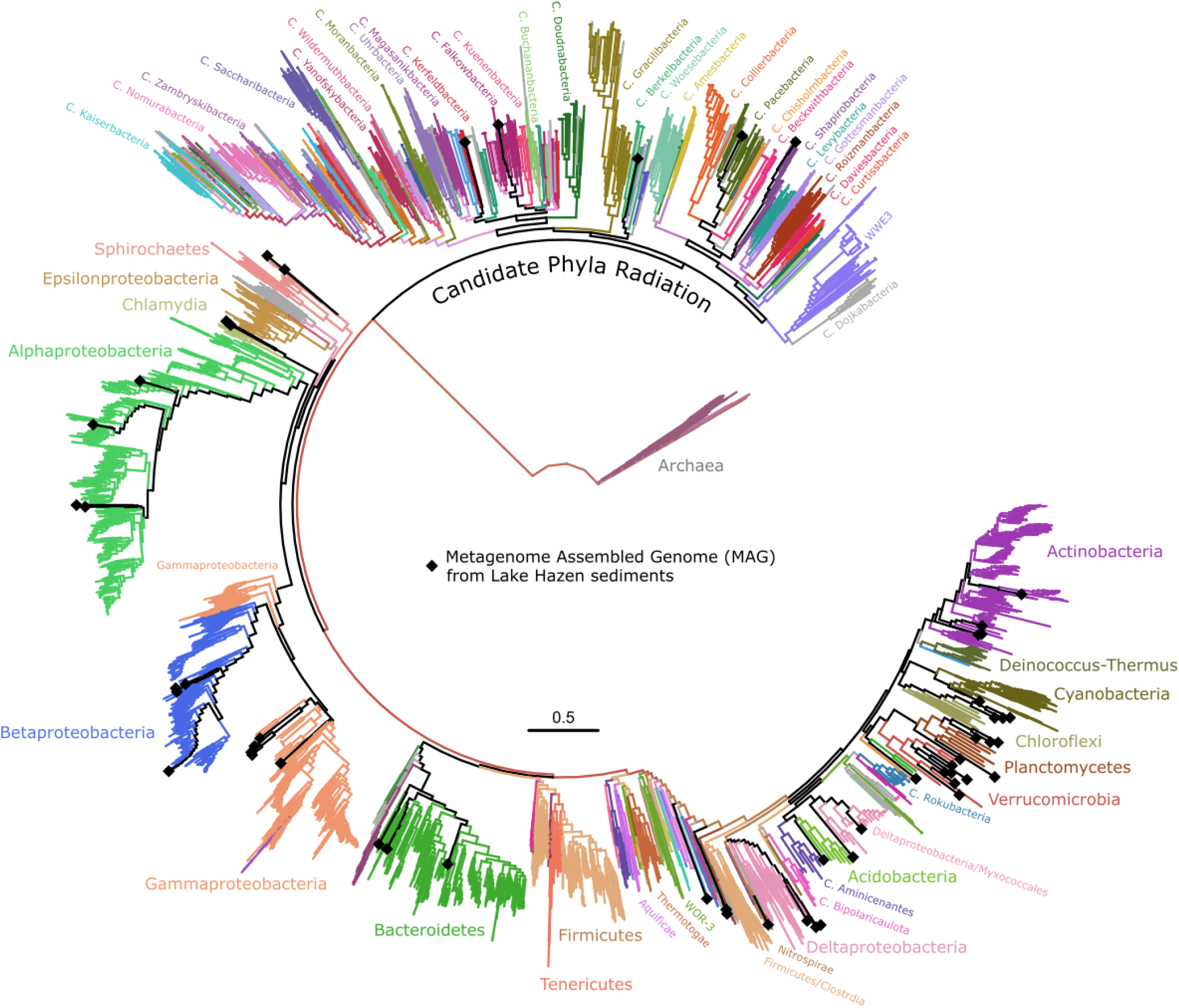
Phylogenetic tree of MAGs aligned against reference genomes annotated at phylum-level based on the NCBI taxonomy database. Genomes identified in NCBI as belonging to each phylum are indicated with different colors, which are matching those in Figure 1. Black diamonds indicate the phylogenetic placement of the MAG assembled in the current study. Scale bar are in unit of number of substitutions per site. Full tree with SH-like node support values is included as supplementary files (Full_tree_with_supports.pdf and Full_tree_with_supports.nwk).

### Lake Hazen MAGs are enriched in genes to strive at low temperatures and oligotrophy

To test if these reconstructed genomes from Lake Hazen possess unique metabolic features, we quantified the presence of select pathways in both the MAGs and a set of reference genomes at the same taxonomic rank (phylum level; *n* = 2,486 genomes), and compared their prevalence (Figure 3). In particular, we found that the marker genes and pathways for cellular metabolism and nutrient cycling (Table S5) were significantly different (Pearson’s χ^2^; *X*^2^ = 361.25; *P* = 0.0001 based on 10,000 permutations).

**Figure 3.**
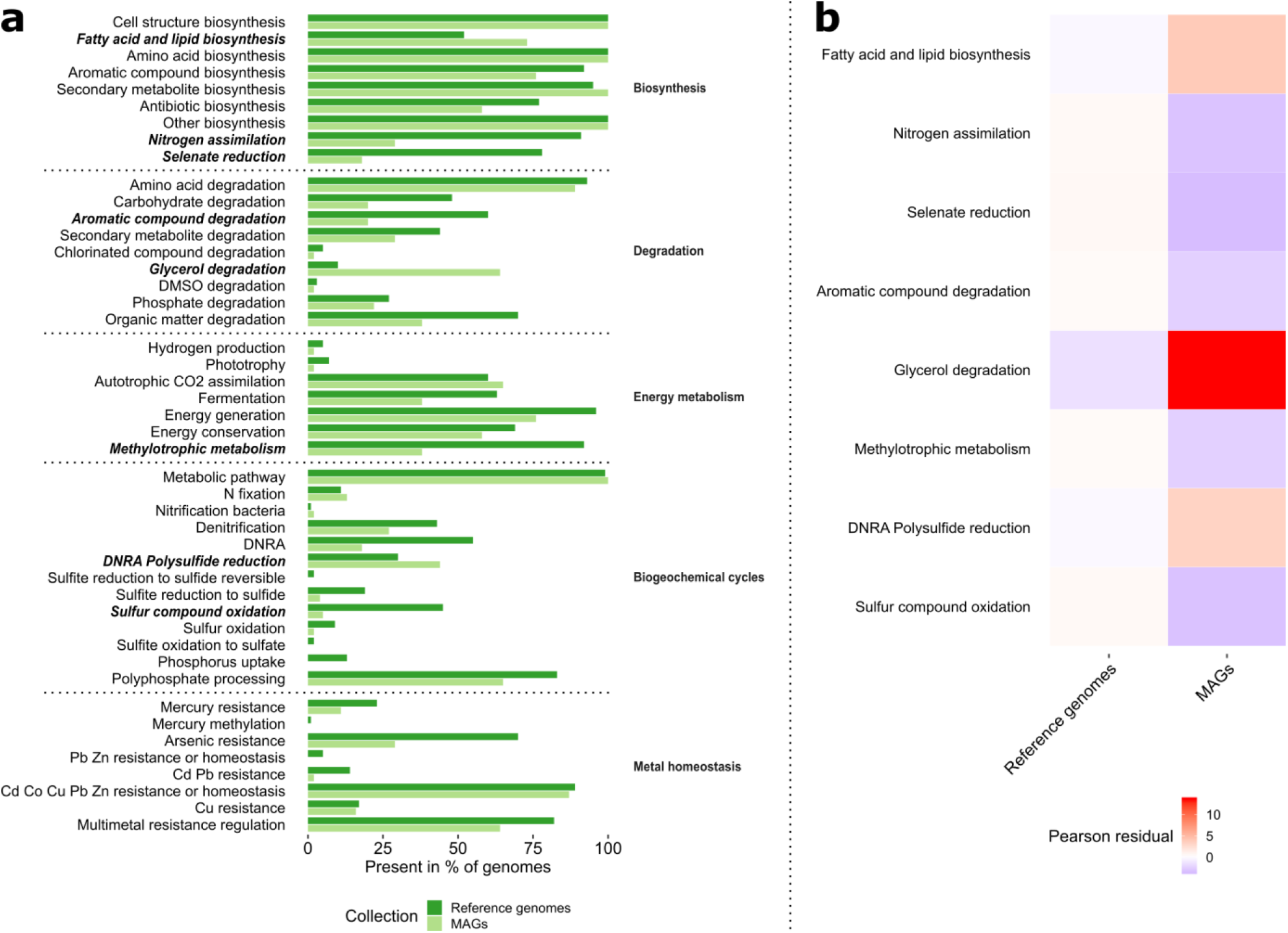
Cellular processes and metabolism in reference genomes and MAGs. (**a**) Percent representation of genomes in each data set (reference genomes and MAGs) with at least one marker pathway or gene for each process or metabolism. Names of pathways or processes with the most differentially abundant presence in MAGs compared to the reference genomes subset to common phyla are indicated in bold italics. (**b**) The most differentially abundant processes according to the Pearson’s χ^2^ test (*P* = 0.001) as a heatmap of their Pearson residual scores.

Among the most different pathways, three were overrepresented in the MAGs (Figure 3b). Among those three, glycerol (glycerophosphodiester) degradation and fatty acid / lipid biosynthesis were more prevalent in the MAGs (Figure 3b). This is likely linked with temperature tolerance and energy conservation strategies. Indeed, both cold stress (Chintalapati et al., 2004) and starvation (Lever et al., 2015) can induce changes in the lipid composition of microbial cell membranes. A high prevalence of fatty acid desaturases was also recently seen in several Antarctic lake metagenomes (Koo et al., 2018), suggesting that these pathways are important for psychrotrophic microbes – given that water temperatures are around 3.5°C below a depth of 50 m (St. Pierre et al., 2019). Alternatively, triacylglycerols could be utilized for energy storage in the form of lipid droplets (Alvarez and Steinbüchel, 2002). In addition to energy storage, lipids, in particular when present as droplets, might also play a role in regulating the stress response in bacteria (Zhang et al., 2017). Both of these roles could be beneficial to the bacteria harboring them in the Lake Hazen sediments, as most nutrients and oxygen are delivered to the lake in glacial meltwater during summer (St. Pierre et al., 2019). This short period of higher nutrient and oxygen availability correlates with a temporary jump in microbial activity (St. Pierre et al., 2019), whose energy stores might be triacylglycerols that are then gradually released to maintain metabolism during the long winter.

The third overrepresented pathway, Dissimilatory Nitrite Reduction to Ammonia (DNRA) / Polysulfide reduction, shows that the MAGs are also enriched in membrane-linked molybdopterin oxidoreductases genes of the NrfD/PsrC family that includes genes coding for tetrathionate-, dimethyl sulfoxide-, polysulfide-, and nitrite reductases (Jormakka et al., 2008). It is more likely that these genes in MAGs (matching the annotation Pfam 03916) are related to sulfur reduction than DNRA, because the more specific markers for DNRA, such as *nrfH*, *nirB* and *nirD*, were rarer in the MAGs than in the reference genomes (Figure 3a). Furthermore, the same marker gene (*nrfD*) has been found for instance in metagenomes from sediments of saline (Ferrer et al., 2011) and freshwater lakes (Lin et al., 2011), and associated there with anaerobic respiration with oxidized sulfur compounds. The *nrfD* gene is thus likely important also for anaerobic respiration in the mostly anoxic sediments of Lake Hazen.

Underrepresented pathways in the Lake Hazen sediment MAGs (Figure 3b) might be underutilized and thus subject to genome streamlining, which is common in oligotrophic organisms (Giovannoni et al., 2014). These pathways include, for example, nitrogen assimilation as NO_3_^−^ and NH_4_^+^. Likely, because of the low concentration of inorganic nitrogen in Lake Hazen (St. Pierre et al., 2019) it might be directly assimilated only by a small number of organisms. The majority of microbes might instead resort to recycling of existing organic nitrogen compounds or nitrogen fixation to fulfill their needs for this nutrient (Figure 3). Similarly, aromatic compound degradation, methylotrophic metabolism and sulfur compound oxidation (largely desulfonation) were rarer in the MAGs than reference genomes. The lower prevalence of these pathways might also be a consequence of low nutrient and substrate availability in Lake Hazen and the low number of methanogens (this study, Emmerton et al., 2016; Ruuskanen et al., 2018; St. Pierre et al., 2019b). In addition, selenate reduction was underrepresented in the MAGs. This pathway is usually important for anaerobic metabolism (Staicu and Barton, 2017), but in Lake Hazen its lower prevalence could similarly reflect the low availability of selenium because the site is distant from anthropogenic sources (e.g., Chapman et al., 2010). It should be noted, that marker genes used to identify the aforementioned pathways may also have been missed due to technological biases, on our focusing the analysis on only the 55 most complete MAGs, and / or their incompleteness. Despite these potential biases however, the underrepresentation of these specific pathways in the MAGs makes sense in the light of the conditions in Lake Hazen sediments. Indeed, Lake Hazen is rapidly changing with the increased delivery of sediment, nutrients, organic carbon and contaminants, perpetuated by enhanced glacial melt throughout the watershed (Lehnherr et al., 2018). The dense and turbid glacial waters entering Lake Hazen flow directly to the bottom of the lake (St. Pierre et al., 2019), and this might effectively lead to the disappearance of oligotrophic ecological niches in the sediments with increased productivity.

### Spatial homogeneity of nutrient / toxic metal cycling in Lake Hazen sediments

Finally, to investigate more closely nutrient and toxic metal cycling in the sediments and evaluate the possible contribution of different microbes to these cycles, we identified marker genes or pathways for these relevant processes. However, because the most parsimonious pathways are calculated using marker genes present in a single genome bin, it is possible that low quality bins could have strongly biased pathway inference (Ye and Doak, 2009). To alleviate this issue, we analyzed differences in abundances of all individual marker genes found in all assembled contigs in the metagenome longer than 1 kbp – and not only the 55 MAGs – using gene-level normalized read coverages summarized per each marker gene (Figure 4).

**Figure 4.**
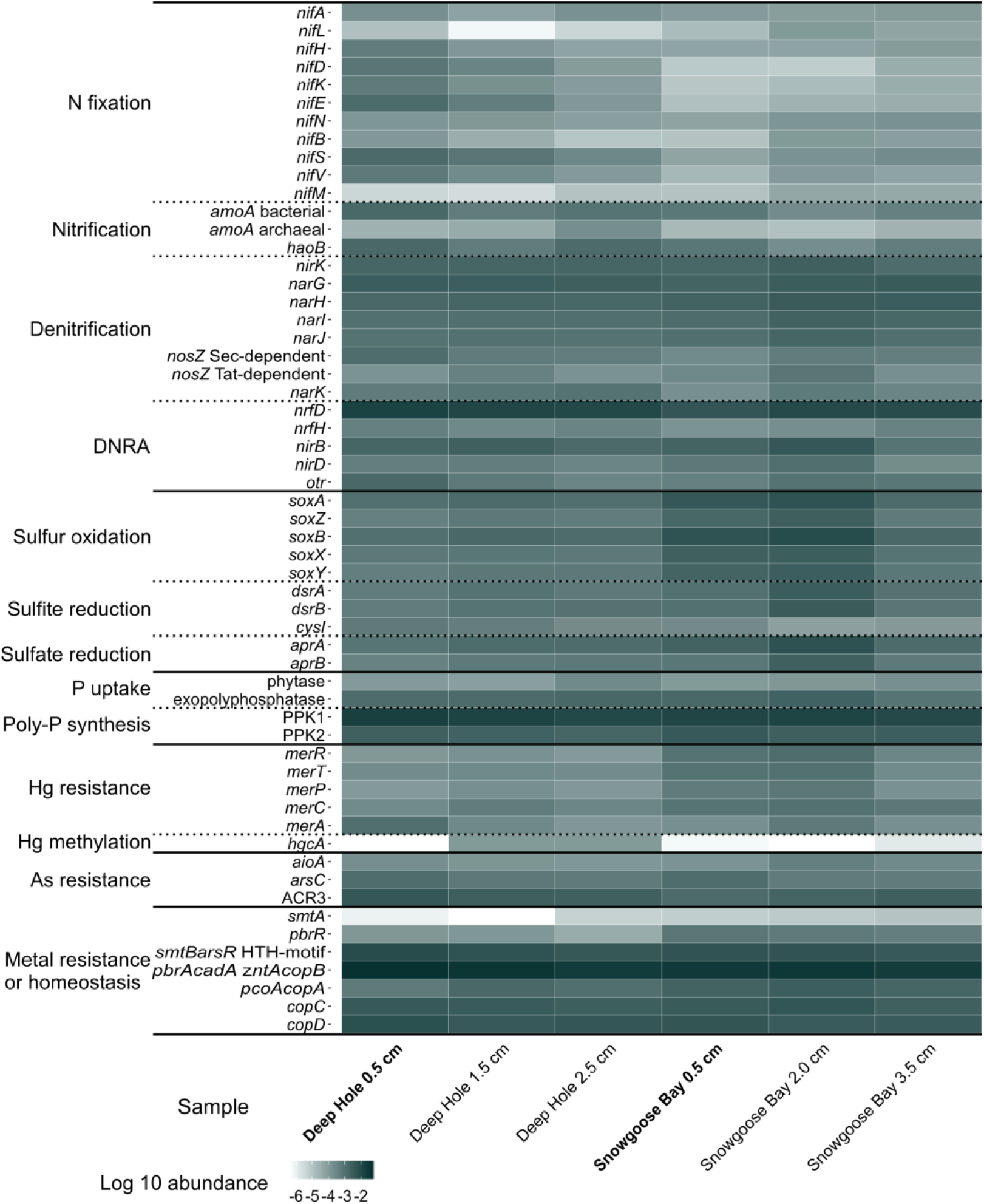
Normalized read coverages of individual marker genes from the whole metagenomic data across the samples. Samples from oxic environments (O_2_ concentration > 0.0 mg L^−1^) are indicated in bold. Differences in abundances of individual marker genes between the two sites (Deep Hole and Snowgoose Bay) and oxygen levels (oxic and anoxic) were not significant (pairwise *t*-test; all *P* > 0.8 after FDR correction). Gene family/domain annotations used to identify the individual marker genes are shown in Table S5.

Comparing within and between sites, as well as oxic (surface samples; < 0.5 cm) vs. anoxic samples (deeper samples are completely anoxic at both sites (Figure S3), we found that none of the individual marker genes for the nutrient cycles were differentially abundant between the two sites (pairwise *t*-tests, all *P* > 0.9 after FDR correction) or between the oxic and anoxic sediment horizons (pairwise *t*-tests, all *P* > 0.8 after FDR correction). Thus, for the next analysis we averaged the normalized abundances of the 55 MAGs across all six samples (the two sites and three depths) to identify the likely key organisms in the nitrogen, sulfur, and mercury cycles – which improved our ability to investigate these more fine-grain patterns of nutrient cycling.

Based on these average read coverages, we identified the most abundant MAGs that included markers for nitrogen, sulfur and mercury cycling (Table S6), as summarized in Table S7 for each ecologically important process in the nutrient cycles and shown in Figure 5. Proteobacteria (mostly beta- and alpha-) were overall the most abundant phylum that had marker genes or pathways for nitrogen and sulfur cycling, but other phyla were often the most abundant when looking at individual processes (SI text; Table S7). The metagenome also contained markers for the full sulfur cycle, although the genes for dissimilatory sulfate reduction to sulfite (*aprAB*) and dissimilatory sulfite reduction to sulfide (*dsrAB*) were not detected specifically in the 55 MAGs (SI text; Figure 5b). While no marker genes for mercury methylation (*hgcA*; Pfam 3599) were found in the MAGs (but were present in 32 reference genomes and 27 low quality bins from the metagenome), six reconstructed genomes harbored *mer*-operon genes involved in mercury resistance, with Alphaproteobacteria as the most abundant of these (Figure 5c). The nitrogen and sulfur cycles in the Lake Hazen sediments appeared to be closely intertwined, catalyzed by taxonomically diverse lineages. Certain MAGs, such as LH_MA_37_3, likely from *Geobacter*, and LH_MA_65_9, might represent in the lake ecosystem highly important organisms that can fix dissolved nitrogen gas, and are thus capable of returning it back into the biological cycles. Furthermore, other MAGs, such as LH_MA_55_1, might represent microbes that play roles simultaneously in both sulfur and nitrogen cycles (SI text). These results are consistent with our previous work that, based on 16S rRNA gene amplicon data, predicted the presence of key functional groups such as aerobic ammonia oxidizers (LH_MA_61_7, likely from Nitrosomonadales), nitrate reducers (e.g., LH_MA_28_10, likely from *Rhodoferax*) and sulfate reducers (presence of *aprAB* genes in the metagenomic dataset) in the Lake Hazen sediments (Ruuskanen et al., 2018).

**Figure 5.**
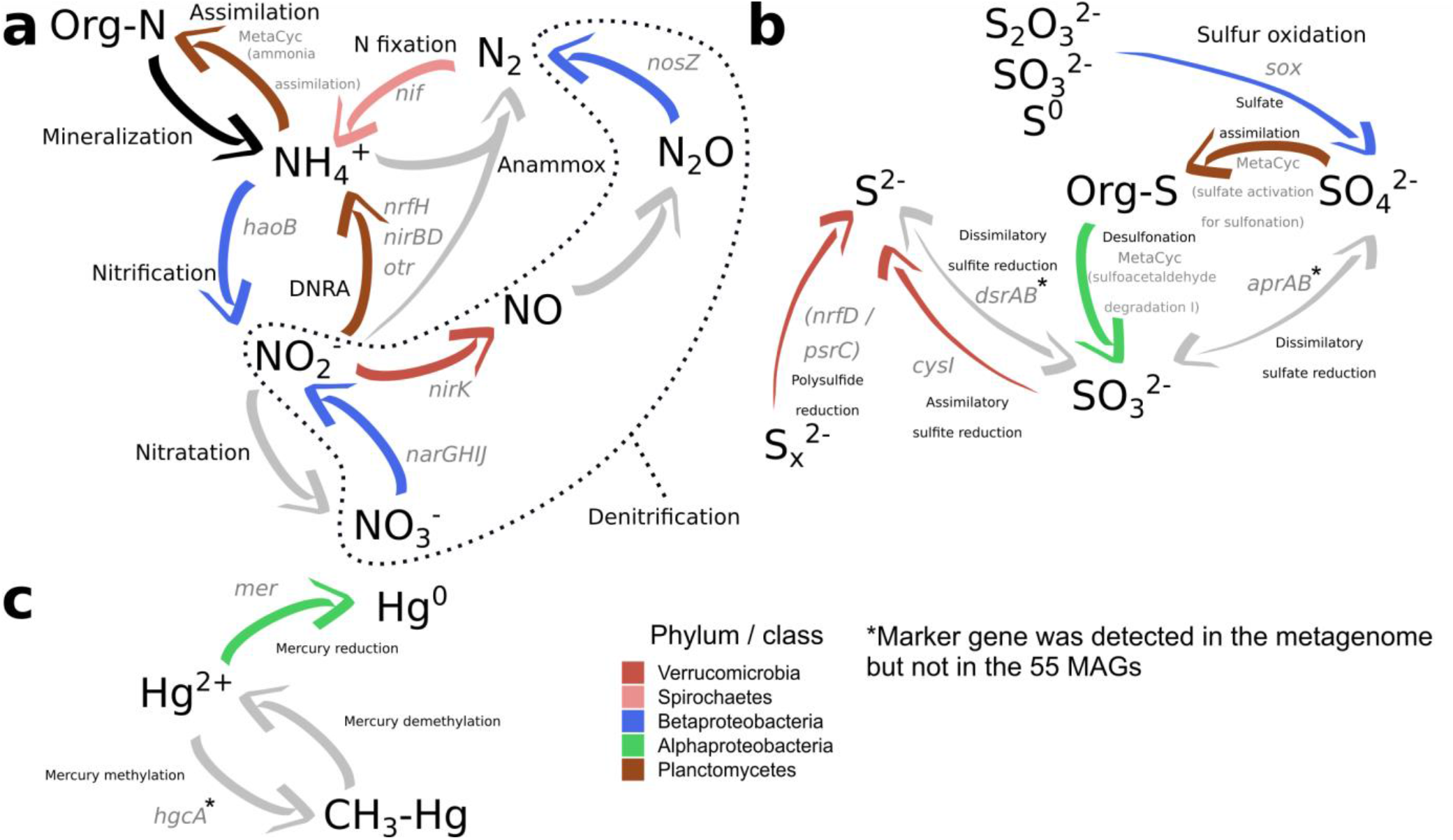
Biogeochemical cycling in the Lake Hazen sediment. Most common phyla participating in each process are indicated. The phylum Proteobacteria is here subdivided into classes. Marker gene symbols or ‘MetaCyc’ for marker pathways are indicated in gray text next to each process. Gray arrows denote processes that were *not* found in the 55 MAGs. If the gene is not indicated, a specific marker for the process could not be identified in the annotation. (**a**) Nitrogen cycle. DNRA: Dissimilatory Reduction of Nitrite to Ammonia. (**b**) Sulfur cycle. (**c**) Mercury cycle.

## Conclusions

We show that in addition to being unique in its location (81°N), dimensions, and volume, Lake Hazen hosts a phylogenetically diverse set of microbes whose reconstructed genomes contain a high prevalence of pathways that make them fit for thriving in a cold (~3.5°C) and oligotrophic environment. This diversity includes organisms from recently discovered groups, such as the Candidate Phyla Radiation, and from uncultured branches of the tree of life. Because metagenomic surveys from arctic lake sediments are currently scarce, our results provide the scientific community with a compositional baseline and functional potential of these ecosystems. Indeed, as the oligotrophic niches that extant microbes inhabit are likely to be affected by ongoing environmental changes in Lake Hazen, changes in the microbial community at the base of the lake ecosystem might have unforeseen consequences, with possible repercussions leading even to higher trophic levels.

## Supporting information

Full_tree_with_supports.pdf

## Acknowledgements

We would like to thank the Poulain and Aris-Brosou lab members at University of Ottawa, and members of the Virta and Hultman labs at University of Helsinki for their helpful comments during data analysis and writing of the manuscript. This work was funded by the Natural Resources Canada / Polar Continental Shelf Project (VStL, AJP), the Natural Sciences and Engineering Research Council of Canada (VStL, SAB, AJP), the ArcticNet Centre for Excellence (VStL, AJP), the Canadian Foundation for Innovation (SAB, AJP) and the Finnish Academy of Science and Letters / the Vilho, Yrjö, and Kalle Väisälä Fund (MR).

## Author contributions

MR wrote the manuscript and analyzed the data. GC participated in sampling, performed the DNA extractions and quality control in the laboratory. KStP and VStL performed the sampling and measured the physicochemical data. AJP and SAB supervised the project. All authors contributed to the writing and accepted the final version of the manuscript.

## References

Abascal, F., Zardoya, R., and Telford, M. J. (2010). TranslatorX: multiple alignment of nucleotide sequences guided by amino acid translations. Nucl. Acids Res., gkq291. doi:10.1093/nar/gkq291.

Alneberg, J., Bjarnason, B. S., de Bruijn, I., Schirmer, M., Quick, J., Ijaz, U. Z., et al. (2014). Binning metagenomic contigs by coverage and composition. Nature Methods 11, 1144–1146. doi:10.1038/nmeth.3103.

Alvarez, H. M., and Steinbüchel, A. (2002). Triacylglycerols in prokaryotic microorganisms. Appl. Microbiol. Biotechnol. 60, 367–376. doi:10.1007/s00253-002-1135-0.

Aris-Brosou, S., and Rodrigue, N. (2012). “The essentials of computational molecular evolution,” in Evolutionary Genomics Methods in Molecular Biology. (Humana Press, Totowa, NJ), 111–152. doi:10.1007/978-1-61779-582-4_4.

Ashburner, M., Ball, C. A., Blake, J. A., Botstein, D., Butler, H., Cherry, J. M., et al. (2000). Gene ontology: tool for the unification of biology. The Gene Ontology Consortium. Nat. Genet. 25, 25–29. doi:10.1038/75556.

Benson, D. A., Cavanaugh, M., Clark, K., Karsch-Mizrachi, I., Lipman, D. J., Ostell, J., et al. (2012). GenBank. Nucleic Acids Research 41, D36–D42. doi:10.1093/nar/gks1195.

Bolger, A. M., Lohse, M., and Usadel, B. (2014). Trimmomatic: a flexible trimmer for Illumina sequence data. Bioinformatics 30, 2114–2120. doi:10.1093/bioinformatics/btu170.

Bowers, R. M., Kyrpides, N. C., Stepanauskas, R., Harmon-Smith, M., Doud, D., Reddy, T. B. K., et al. (2017). Minimum information about a single amplified genome (MISAG) and a metagenome-assembled genome (MIMAG) of bacteria and archaea. Nature Biotechnology 35, 725–731. doi:10.1038/nbt.3893.

Brown, C. T., Hug, L. A., Thomas, B. C., Sharon, I., Castelle, C. J., Singh, A., et al. (2015). Unusual biology across a group comprising more than 15% of domain Bacteria. Nature 523, 208.

Buchfink, B., Xie, C., and Huson, D. H. (2015). Fast and sensitive protein alignment using DIAMOND. Nature Methods 12, 59–60. doi:10.1038/nmeth.3176.

Campbell, J. H., O’Donoghue, P., Campbell, A. G., Schwientek, P., Sczyrba, A., Woyke, T., et al. (2013). UGA is an additional glycine codon in uncultured SR1 bacteria from the human microbiota. PNAS 110, 5540–5545. doi:10.1073/pnas.1303090110.

Campello, R. J. G. B., Moulavi, D., and Sander, J. (2013). Density-based clustering based on hierarchical density estimates. in Advances in Knowledge Discovery and Data Mining Lecture Notes in Computer Science. (Springer, Berlin, Heidelberg), 160–172. doi:10.1007/978-3-642-37456-2_14.

Capella-Gutiérrez, S., Silla-Martínez, J. M., and Gabaldón, T. (2009). trimAl: a tool for automated alignment trimming in large-scale phylogenetic analyses. Bioinformatics 25, 1972–1973. doi:10.1093/bioinformatics/btp348.

Carr, S. A., Schubotz, F., Dunbar, R. B., Mills, C. T., Dias, R., Summons, R. E., et al. (2018). Acetoclastic *Methanosaeta* are dominant methanogens in organic-rich Antarctic marine sediments. The ISME Journal 12, 330–342. doi:10.1038/ismej.2017.150.

Caspi, R., Billington, R., Fulcher, C. A., Keseler, I. M., Kothari, A., Krummenacker, M., et al. (2018). The MetaCyc database of metabolic pathways and enzymes. Nucleic Acids Res 46, D633–D639. doi:10.1093/nar/gkx935.

Chapman, P. M., Adams, W. J., Brooks, M., Delos, C. G., Luoma, S. N., Maher, W. A., et al. (2010). Ecological Assessment of Selenium in the Aquatic Environment. CRC Press.

Chintalapati, S., Kiran, M. D., and Shivaji, S. (2004). Role of membrane lipid fatty acids in cold adaptation. Cell. Mol. Biol. (Noisy-le-grand) 50, 631–642.

Crump, B. C., Amaral-Zettler, L. A., and Kling, G. W. (2012). Microbial diversity in arctic freshwaters is structured by inoculation of microbes from soils. ISME J 6, 1629–1639. doi:10.1038/ismej.2012.9.

Danczak, R. E., Johnston, M. D., Kenah, C., Slattery, M., Wrighton, K. C., and Wilkins, M. J. (2017). Members of the Candidate Phyla Radiation are functionally differentiated by carbon- and nitrogen-cycling capabilities. Microbiome 5. doi:10.1186/s40168-017-0331-1.

Delmont, T. O., Quince, C., Shaiber, A., Esen, Ö. C., Lee, S. T., Rappé, M. S., et al. (2018). Nitrogen-fixing populations of Planctomycetes and Proteobacteria are abundant in surface ocean metagenomes. Nat Microbiol 3, 804–813. doi:10.1038/s41564-018-0176-9.

Eddy, S. R. (2009). A new generation of homology search tools based on probabilistic inference. Genome Inform 23, 205–211.

Emmerton, C. A., St. Louis, V. L., Lehnherr, I., Graydon, J. A., Kirk, J. L., and Rondeau, K. J. (2016). The importance of freshwater systems to the net atmospheric exchange of carbon dioxide and methane with a rapidly changing high Arctic watershed. Biogeosciences 13, 5849–5863. doi:10.5194/bg-13-5849-2016.

Eren, A. M., Esen, Ö. C., Quince, C., Vineis, J. H., Morrison, H. G., Sogin, M. L., et al. (2015). Anvi’o: an advanced analysis and visualization platform for ‘omics data. PeerJ 3, e1319. doi:10.7717/peerj.1319.

Ferrer, M., Guazzaroni, M.-E., Richter, M., García-Salamanca, A., Yarza, P., Suárez-Suárez, A., et al. (2011). Taxonomic and Functional Metagenomic Profiling of the Microbial Community in the Anoxic Sediment of a Sub-saline Shallow Lake (Laguna de Carrizo, Central Spain). Microbial Ecology 62, 824–837.

Filippidou, S., Junier, T., Wunderlin, T., Lo, C.-C., Li, P.-E., Chain, P. S., et al. (2015). Under-detection of endospore-forming Firmicutes in metagenomic data. Computational and Structural Biotechnology Journal 13, 299–306. doi:10.1016/j.csbj.2015.04.002.

Finn, R. D., Bateman, A., Clements, J., Coggill, P., Eberhardt, R. Y., Eddy, S. R., et al. (2014). Pfam: the protein families database. Nucleic Acids Res 42, D222–D230. doi:10.1093/nar/gkt1223.

France, R. L. (1993). The Lake Hazen Trough: A late winter oasis in a polar desert. Biological Conservation 63, 149–151. doi:10.1016/0006-3207(93)90503-S.

Galperin, M. Y., Makarova, K. S., Wolf, Y. I., and Koonin, E. V. (2015). Expanded microbial genome coverage and improved protein family annotation in the COG database. Nucleic Acids Res 43, D261–D269. doi:10.1093/nar/gku1223.

Giovannoni, S. J., Cameron Thrash, J., and Temperton, B. (2014). Implications of streamlining theory for microbial ecology. ISME J 8, 1553–1565. doi:10.1038/ismej.2014.60.

Hahsler, M., Piekenbrock, M., Arya, S., and Mount, D. (2018). dbscan: Density Based Clustering of Applications with Noise (DBSCAN) and Related Algorithms. Available at: https://CRAN.R-project.org/package=dbscan [Accessed June 4, 2018].

Howe, A. C., Jansson, J. K., Malfatti, S. A., Tringe, S. G., Tiedje, J. M., and Brown, C. T. (2014). Tackling soil diversity with the assembly of large, complex metagenomes. PNAS, 201402564. doi:10.1073/pnas.1402564111.

Hug, L. A., Baker, B. J., Anantharaman, K., Brown, C. T., Probst, A. J., Castelle, C. J., et al. (2016). A new view of the tree of life. Nature Microbiology 1, 16048.

Hunter, S., Jones, P., Mitchell, A., Apweiler, R., Attwood, T. K., Bateman, A., et al. (2012). InterPro in 2011: new developments in the family and domain prediction database. Nucleic Acids Res 40, D306–D312. doi:10.1093/nar/gkr948.

Hyatt, D., Chen, G.-L., LoCascio, P. F., Land, M. L., Larimer, F. W., and Hauser, L. J. (2010). Prodigal: prokaryotic gene recognition and translation initiation site identification. BMC Bioinformatics 11, 119. doi:10.1186/1471-2105-11-119.

IPCC (2018). Global warming of 1.5 C An IPCC Special Report on the impacts of global warming of 1.5 C above pre-industrial levels and related global greenhouse gas emission pathways, in the context of strengthening the global response to the threat of climate change, sustainable development, and efforts to eradicate poverty., eds. V. Masson-Delmotte, P. Zhai, H. O. Pörtner, D. Roberts, J. Skea, P. R. Shukla, et al. Summary for Policymakers Edited by Science Officer Science Assistant ….

Jones, P., Binns, D., Chang, H.-Y., Fraser, M., Li, W., McAnulla, C., et al. (2014). InterProScan 5: genome-scale protein function classification. Bioinformatics 30, 1236–1240. doi:10.1093/bioinformatics/btu031.

Jormakka, M., Yokoyama, K., Yano, T., Tamakoshi, M., Akimoto, S., Shimamura, T., et al. (2008). Molecular mechanism of energy conservation in polysulfide respiration. Nat Struct Mol Biol 15, 730–737. doi:10.1038/nsmb.1434.

Kanehisa, M., Furumichi, M., Tanabe, M., Sato, Y., and Morishima, K. (2017). KEGG: new perspectives on genomes, pathways, diseases and drugs. Nucleic Acids Res 45, D353–D361. doi:10.1093/nar/gkw1092.

Katoh, K., and Standley, D. M. (2013). MAFFT Multiple Sequence Alignment Software Version 7: Improvements in Performance and Usability. Molecular Biology and Evolution 30, 772–780. doi:10.1093/molbev/mst010.

Koo, H., Hakim, J. A., Morrow, C. D., Crowley, M. R., Andersen, D. T., and Bej, A. K. (2018). Metagenomic Analysis of Microbial Community Compositions and Cold-Responsive Stress Genes in Selected Antarctic Lacustrine and Soil Ecosystems. Life (Basel) 8. doi:10.3390/life8030029.

Krijthe, J., and van der Maaten, L. (2017). rtsne: T-distributed stochastic neighbor embedding using a Barnes-Hut implementation. Available at: https://cran.r-project.org/web/packages/Rtsne/index.html [Accessed November 29, 2017].

Langmead, B., and Salzberg, S. L. (2012). Fast gapped-read alignment with Bowtie 2. Nat. Methods 9, 357–359. doi:10.1038/nmeth.1923.

Lau, N.-S., Zarkasi, K. Z., Md Sah, A. S. R., and Shu-Chien, A. C. (2018). Diversity and Coding Potential of the Microbiota in the Photic and Aphotic Zones of Tropical Man-Made Lake with Intensive Aquaculture Activities: a Case Study on Temengor Lake, Malaysia. Microb Ecol. doi:10.1007/s00248-018-1283-0.

Lehnherr, I., Louis, V. L. S., Sharp, M., Gardner, A. S., Smol, J. P., Schiff, S. L., et al. (2018). The world’s largest High Arctic lake responds rapidly to climate warming. Nature Communications 9, 1290. doi:10.1038/s41467-018-03685-z.

León‐Zayas, R., Peoples, L., Biddle, J. F., Podell, S., Novotny, M., Cameron, J., et al. (2017). The metabolic potential of the single cell genomes obtained from the Challenger Deep, Mariana Trench within the candidate superphylum Parcubacteria (OD1). Environmental Microbiology 19, 2769–2784. doi:10.1111/1462-2920.13789.

Lever, M. A., Rogers, K. L., Lloyd, K. G., Overmann, J., Schink, B., Thauer, R. K., et al. (2015). Life under extreme energy limitation: a synthesis of laboratory- and field-based investigations. FEMS Microbiol Rev 39, 688–728. doi:10.1093/femsre/fuv020.

Li, D., Liu, C.-M., Luo, R., Sadakane, K., and Lam, T.-W. (2015). MEGAHIT: an ultra-fast single-node solution for large and complex metagenomics assembly via succinct de Bruijn graph. Bioinformatics 31, 1674–1676. doi:10.1093/bioinformatics/btv033.

Lin, W., Jogler, C., Schüler, D., and Pan, Y. (2011). Metagenomic Analysis Reveals Unexpected Subgenomic Diversity of Magnetotactic Bacteria within the Phylum Nitrospirae. Appl. Environ. Microbiol. 77, 323–326. doi:10.1128/AEM.01476-10.

Louca, S., Parfrey, L. W., and Doebeli, M. (2016). Decoupling function and taxonomy in the global ocean microbiome. Science 353, 1272–1277. doi:10.1126/science.aaf4507.

Mai, U., and Mirarab, S. (2018). TreeShrink: fast and accurate detection of outlier long branches in collections of phylogenetic trees. BMC Genomics 19, 272. doi:10.1186/s12864-018-4620-2.

McMurdie, P. J., and Holmes, S. (2013). phyloseq: An R package for reproducible interactive analysis and graphics of microbiome census data. PLoS ONE 8, e61217. doi:10.1371/journal.pone.0061217.

Milner, A. M., Khamis, K., Battin, T. J., Brittain, J. E., Barrand, N. E., Füreder, L., et al. (2017). Glacier shrinkage driving global changes in downstream systems. PNAS 114, 9770–9778. doi:10.1073/pnas.1619807114.

Mohit, V., Culley, A., Lovejoy, C., Bouchard, F., and Vincent, W. F. (2017). Hidden biofilms in a far northern lake and implications for the changing Arctic. NPJ Biofilms Microbiomes 3. doi:10.1038/s41522-017-0024-3.

Nawrocki, E. P., and Eddy, S. R. (2010). ssu-align: a tool for structural alignment of SSU rRNA sequences. URL http://selab.janelia.org/software.html.

Oksanen, J., Blanchet, F. G., Friendly, M., Kindt, R., Legendre, P., McGlinn, D., et al. (2018). vegan: Community Ecology Package. Available at: https://CRAN.R-project.org/package=vegan [Accessed June 4, 2018].

Paradis, E., Claude, J., and Strimmer, K. (2004). APE: Analyses of phylogenetics and evolution in R language. Bioinformatics 20, 289–290. doi:10.1093/bioinformatics/btg412.

Parks, D. H., Chuvochina, M., Waite, D. W., Rinke, C., Skarshewski, A., Chaumeil, P.-A., et al. (2018). A standardized bacterial taxonomy based on genome phylogeny substantially revises the tree of life. Nature biotechnology.

Parks, D. H., Imelfort, M., Skennerton, C. T., Hugenholtz, P., and Tyson, G. W. (2015). CheckM: assessing the quality of microbial genomes recovered from isolates, single cells, and metagenomes. Genome Res. 25, 1043–1055. doi:10.1101/gr.186072.114.

Pavoine, S., Dufour, A.-B. A.-B., and Chessel, D. (2004). From dissimilarities among species to dissimilarities among communities: a double principal coordinate analysis. J. Theor. Biol. 228, 523–537. doi:10.1016/j.jtbi.2004.02.014.

Pericard, P., Dufresne, Y., Couderc, L., Blanquart, S., Touzet, H., and Birol, I. (2018). MATAM: reconstruction of phylogenetic marker genes from short sequencing reads in metagenomes. Bioinformatics 34, 585–591. doi:10.1093/bioinformatics/btx644.

Poulain, A. J., Aris-Brosou, S., Blais, J. M., Brazeau, M., Keller, W. (Bill), and Paterson, A. M. (2015). Microbial DNA records historical delivery of anthropogenic mercury. The ISME Journal. doi:10.1038/ismej.2015.86.

Pruesse, E., Peplies, J., and Glöckner, F. O. (2012). SINA: Accurate high-throughput multiple sequence alignment of ribosomal RNA genes. Bioinformatics 28, 1823–1829. doi:10.1093/bioinformatics/bts252.

Quast, C., Pruesse, E., Yilmaz, P., Gerken, J., Schweer, T., Yarza, P., et al. (2013). The SILVA ribosomal RNA gene database project: Improved data processing and web-based tools. Nucl. Acids Res. 41, D590–D596. doi:10.1093/nar/gks1219.

R Core Team (2018). R: A language and environment for statistical computing. Vienna, Austria: R Foundation for Statistical Computing Available at: https://www.R-project.org/.

Rautio, M., Dufresne, F., Laurion, I., Bonilla, S., Vincent, W. F., and Christoffersen, K. S. (2011). Shallow freshwater ecosystems of the circumpolar Arctic1. Écoscience; Sainte-Foy 18, 204–222.

Rinke, C., Schwientek, P., Sczyrba, A., Ivanova, N. N., Anderson, I. J., Cheng, J.-F., et al. (2013). Insights into the phylogeny and coding potential of microbial dark matter. Nature 499, 431–437. doi:10.1038/nature12352.

Romanovsky, V. E., Smith, S. L., and Christiansen, H. H. (2010). Permafrost thermal state in the polar Northern Hemisphere during the international polar year 2007–2009: a synthesis. Permafrost and Periglacial Processes 21, 106–116. doi:10.1002/ppp.689.

Ruuskanen, M. O., St. Pierre, K. A., St. Louis, V. L., Aris-Brosou, S., and Poulain, A. J. (2018). Physicochemical Drivers of Microbial Community Structure in Sediments of Lake Hazen, Nunavut, Canada. Front. Microbiol. 9. doi:10.3389/fmicb.2018.01138.

Selengut, J. D., Haft, D. H., Davidsen, T., Ganapathy, A., Gwinn-Giglio, M., Nelson, W. C., et al. (2007). TIGRFAMs and Genome Properties: tools for the assignment of molecular function and biological process in prokaryotic genomes. Nucleic Acids Res. 35, D260–264. doi:10.1093/nar/gkl1043.

Shokralla, S., Spall, J. L., Gibson, J. F., and Hajibabaei, M. (2012). Next-generation sequencing technologies for environmental DNA research. Molecular Ecology 21, 1794–1805. doi:10.1111/j.1365-294X.2012.05538.x.

Soper, J. H., and Powell, J. M. (1985). Botanical studies in the Lake Hazen Region, northern Ellesmere Island, Northwest Territories, Canada.

St. Pierre, K. A., St. Louis, V. L., Lehnherr, I., Schiff, S. L., Muir, D. C. G., Poulain, A. J., et al. (2019). Contemporary limnology of the rapidly changing glacierized watershed of the world’s largest High Arctic lake. Scientific Reports. doi:10.1038/s41598-019-39918-4.

Staicu, L. C., and Barton, L. L. (2017). “Bacterial Metabolism of Selenium—For Survival or Profit,” in Bioremediation of Selenium Contaminated Wastewater, ed. E. D. van Hullebusch (Cham: Springer International Publishing), 1–31. doi:10.1007/978-3-319-57831-6_1.

Stoeva, M. K., Aris-Brosou, S., Chételat, J., Hintelmann, H., Pelletier, P., and Poulain, A. J. (2014). Microbial community structure in lake and wetland sediments from a High Arctic polar desert revealed by targeted transcriptomics. PLOS ONE 9, e89531. doi:10.1371/journal.pone.0089531.

Suzuki, M. T., and Giovannoni, S. J. (1996). Bias caused by template annealing in the amplification of mixtures of 16S rRNA genes by PCR. Appl. Environ. Microbiol. 62, 625–630.

Tatusov, R. L., Fedorova, N. D., Jackson, J. D., Jacobs, A. R., Kiryutin, B., Koonin, E. V., et al. (2003). The COG database: an updated version includes eukaryotes. BMC Bioinformatics 4, 41. doi:10.1186/1471-2105-4-41.

The Gene Ontology Consortium (2017). Expansion of the Gene Ontology knowledgebase and resources. Nucleic Acids Res. 45, D331–D338. doi:10.1093/nar/gkw1108.

van der Maaten, L., and Hinton, G. (2008). Visualizing data using t-SNE. Journal of Machine Learning Research 9, 2579–2605.

Vavourakis, C. D., Andrei, A.-S., Mehrshad, M., Ghai, R., Sorokin, D. Y., and Muyzer, G. (2018). A metagenomics roadmap to the uncultured genome diversity in hypersaline soda lake sediments. Microbiome 6, 168. doi:10.1186/s40168-018-0548-7.

Vigneron, A., Lovejoy, C., Cruaud, P., Kalenitchenko, D., Culley, A., and Vincent, W. F. (2019). Contrasting Winter Versus Summer Microbial Communities and Metabolic Functions in a Permafrost Thaw Lake. Front. Microbiol. 10. doi:10.3389/fmicb.2019.01656.

Wang, N. F., Zhang, T., Yang, X., Wang, S., Yu, Y., Dong, L. L., et al. (2016). Diversity and composition of bacterial community in soils and lake sediments from an arctic lake area. Front Microbiol 7. doi:10.3389/fmicb.2016.01170.

Wang, N., Guo, Y., Li, G., Xia, Y., Ma, M., Zang, J., et al. (2019). Geochemical-Compositional-Functional Changes in Arctic Soil Microbiomes Post Land Submergence Revealed by Metagenomics. Microbes Environ., ME18091. doi:10.1264/jsme2.ME18091.

Wang, Q., Garrity, G. M., Tiedje, J. M., and Cole, J. R. (2007). Naïve Bayesian Classifier for Rapid Assignment of rRNA Sequences into the New Bacterial Taxonomy. Appl. Environ. Microbiol. 73, 5261–5267. doi:10.1128/AEM.00062-07.

Wurzbacher, C., Nilsson, R. H., Rautio, M., and Peura, S. (2017). Poorly known microbial taxa dominate the microbiome of permafrost thaw ponds. ISME J 11, 1938–1941. doi:10.1038/ismej.2017.54.

Ye, Y., and Doak, T. G. (2009). A Parsimony Approach to Biological Pathway Reconstruction/Inference for Genomes and Metagenomes. PLOS Computational Biology 5, e1000465. doi:10.1371/journal.pcbi.1000465.

Yu, G., Smith, D. K., Zhu, H., Guan, Y., and Lam, T. T.-Y. (2017). GGTREE : an R package for visualization and annotation of phylogenetic trees with their covariates and other associated data. Methods in Ecology and Evolution 8, 28–36. doi:10.1111/2041-210X.12628.

Zhang, C., Yang, L., Ding, Y., Wang, Y., Lan, L., Ma, Q., et al. (2017). Bacterial lipid droplets bind to DNA via an intermediary protein that enhances survival under stress. Nature Communications 8, 15979. doi:10.1038/ncomms15979.

Zhang, Z., Schwartz, S., Wagner, L., and Miller, W. (2000). A greedy algorithm for aligning DNA sequences. J. Comput. Biol. 7, 203–214. doi:10.1089/10665270050081478.

Zhou, J., Bruns, M. A., and Tiedje, J. M. (1996). DNA recovery from soils of diverse composition. Appl. Environ. Microbiol. 62, 316–322.

Zhou, J., He, Z., Yang, Y., Deng, Y., Tringe, S. G., and Alvarez-Cohen, L. (2015). High-Throughput Metagenomic Technologies for Complex Microbial Community Analysis: Open and Closed Formats. mBio 6, e02288–14. doi:10.1128/mBio.02288-14.

