## Supplementary material for "Microbial genomes retrieved from High Arctic lake sediments encode for adaptation to cold and oligotrophic environments": SI text

Supplementary tables and the full phylogenetic tree used for MAG taxonomy assignments are provided as separate files.

### 1 Supplementary figures and text

#### 1.1 Sampling

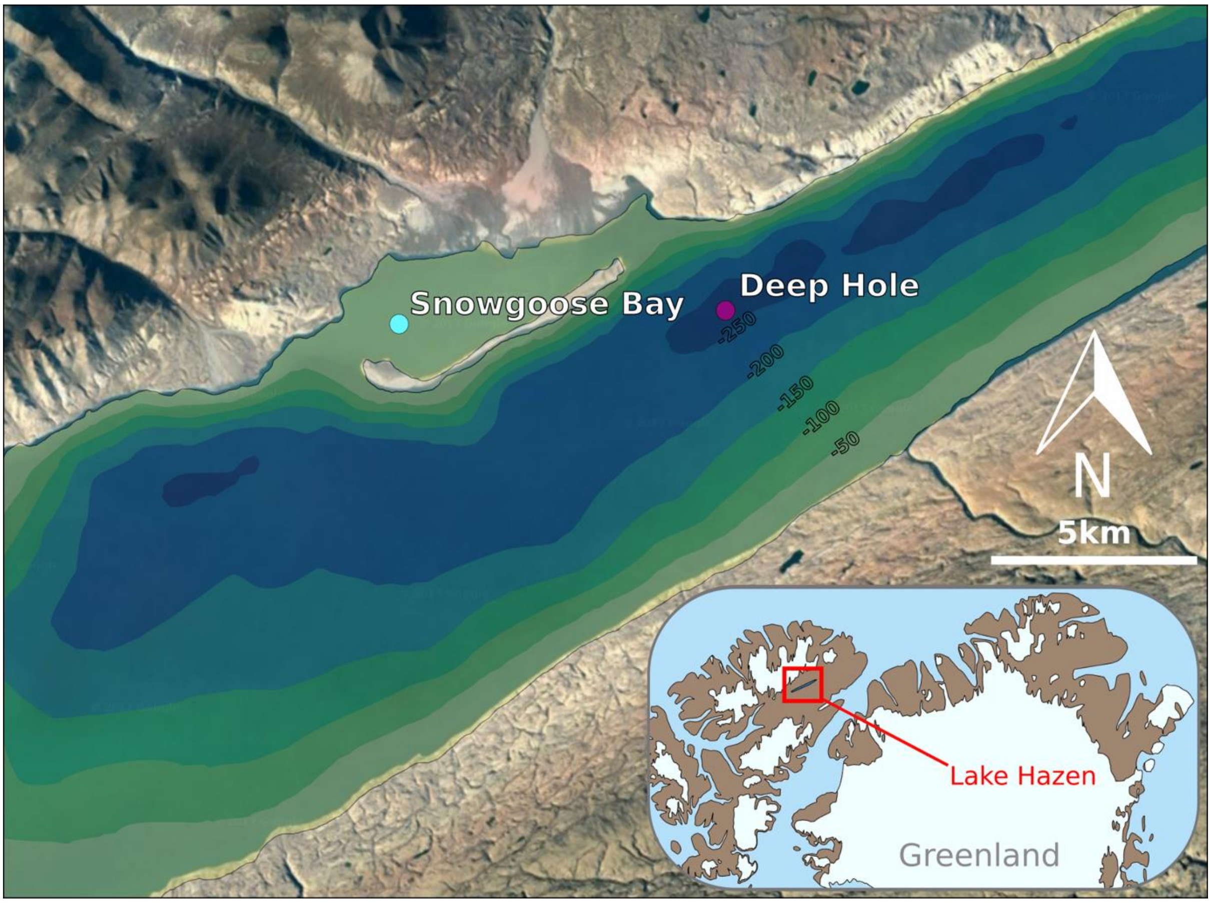

**Figure S1.** Sampling sites at Lake Hazen and location of the site on northern Ellesmere Island, Nunavut, Canada (inlay). Landsat-7 satellite image courtesy of the U.S. Geological Survey. Bathymetric data adapted from Köck et al. (2012).

#### 1.2 DNA extraction and sequencing

PCR for the *glnA* gene to check for DNA quality was performed with the primers GS1*β* (5'-GAT GCC GCC GAT GTA GTA-3') and GS2*γ* (5'-AAG ACC GCG ACC TTY ATG CC-3'), which generate a 153 or 156 bp fragment of the gene (Hurt et al., 2001). Amplifications were performed in 25 µL reaction volumes, each containing 12.5 µL of EconoTaq PLUS GREEN 2X Master Mix (Lucigen Corporation, Middleton, WI, USA), 1 µL of each forward and reverse primer (25 µM), 1 µL of template (DNA extract) and 9.5 µL of $H_2O$. Reaction cycling consisted of initial disassociation at 94ºC for 2 min, followed by disassociation at 94ºC for 30s, annealing at 55ºC for 30s and elongation at 72ºC for 30s for 30 times, with a final elongation at 72ºC for 5 min. Amplification from all DNA extracts, but not the $H_2O$ PCR controls, was confirmed by electrophoresis.

#### 1.3 Assembly

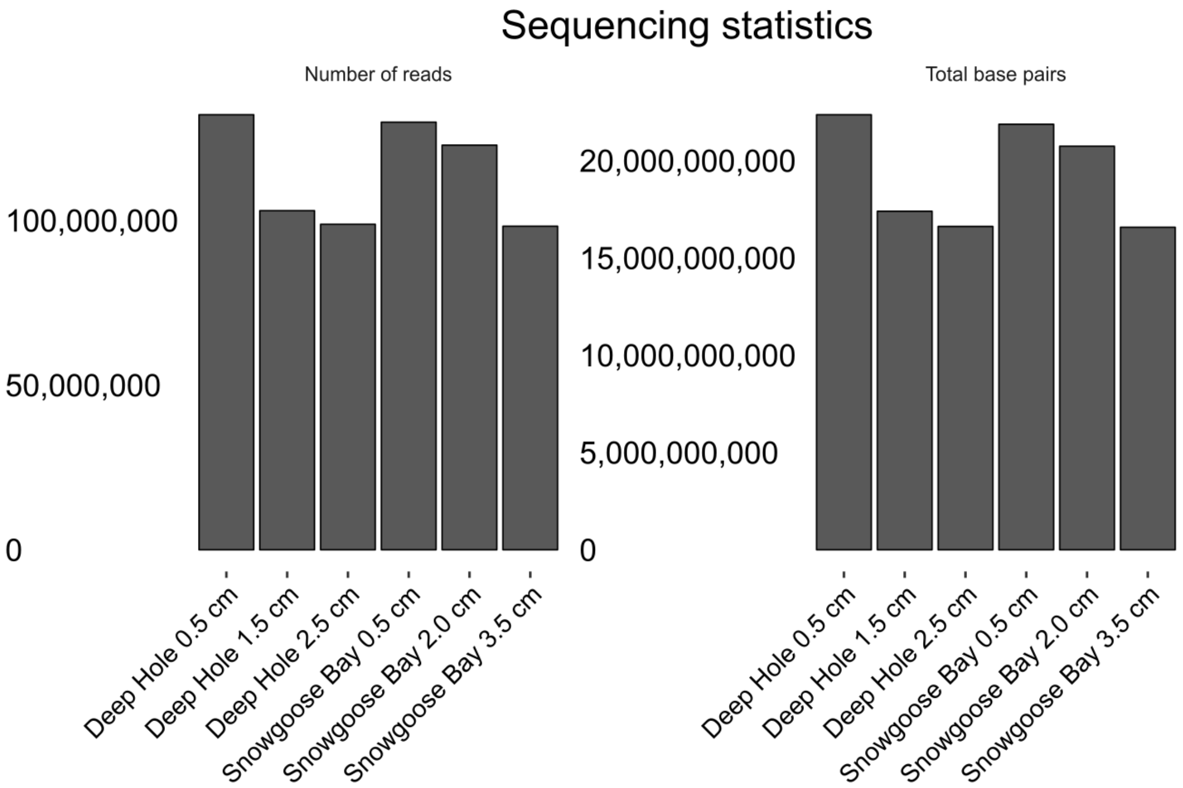

**Figure S2.** Raw number of reads and base pairs per sample.

#### 1.4 Chemistry

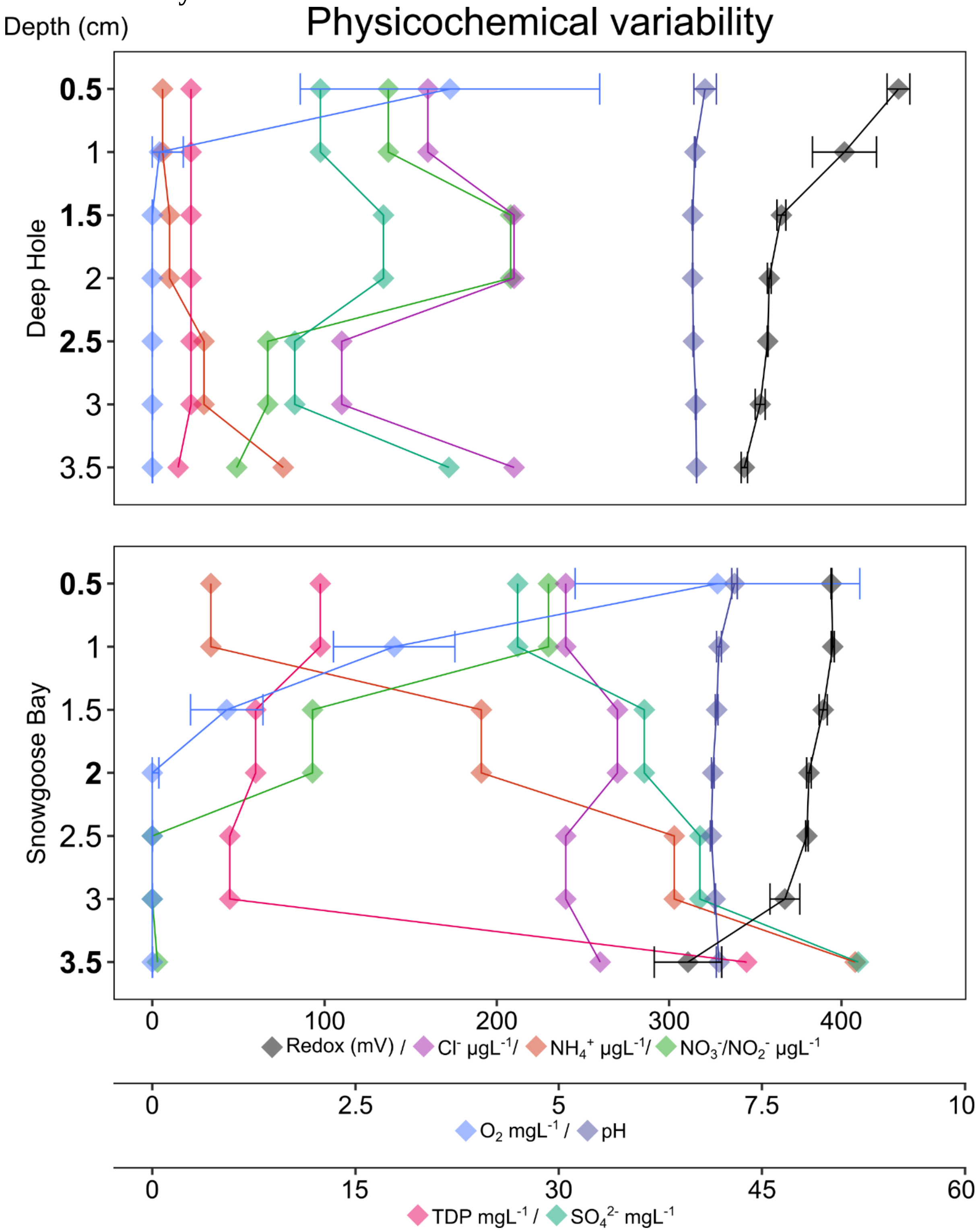

**Figure S3.** Physicochemical variability in the samples from Lake Hazen sediments. Specific scales for measured variables are indicated as three separate x-axes. Sediment depths where DNA was extracted from in each core are indicated with bold font.

#### 1.5 Community structure

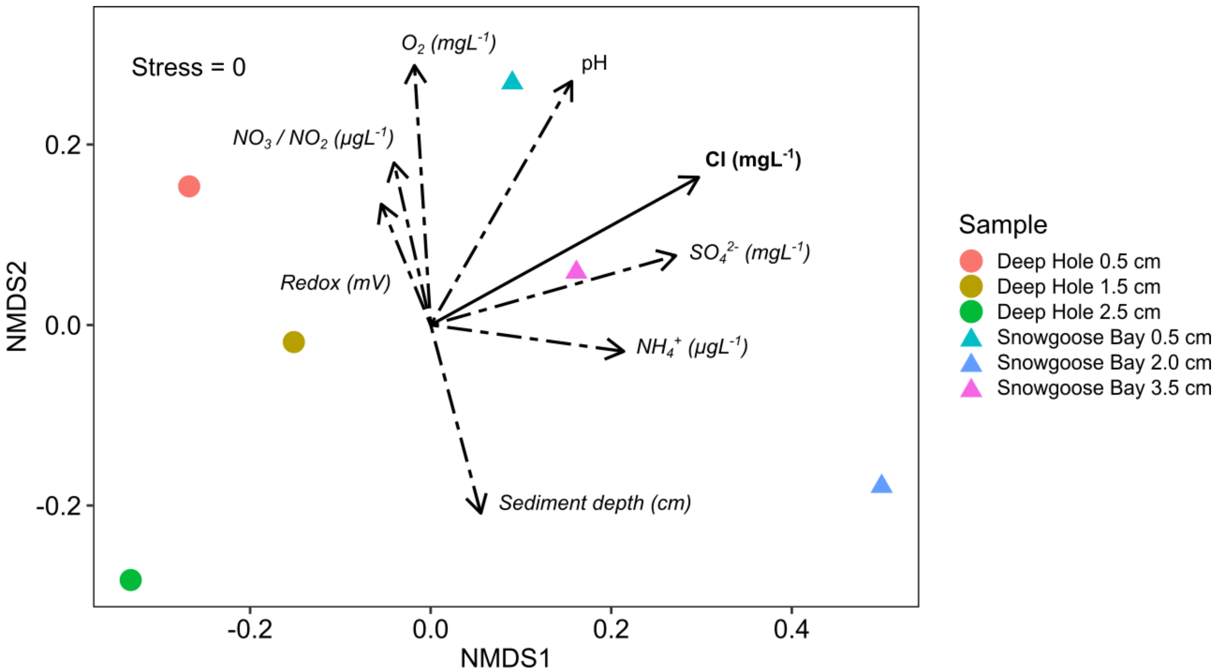

**Figure S4.** NMDS ordination of Bray-Curtis dissimilarity of the microbial communities ($n$ = 166) of the six samples, based on assembly of 16S rRNA fragments. NMDS stress is indicated in the top left corner and vectors for physicochemical variables are overlaid. Continuous variables other than [$Cl^-$] (solid vector; $P = 0.03$) were not significantly linearly correlated with the ordination (dashed vectors; $P > 0.1$), and the samples could not be differentiated by site ($P = 0.1$).

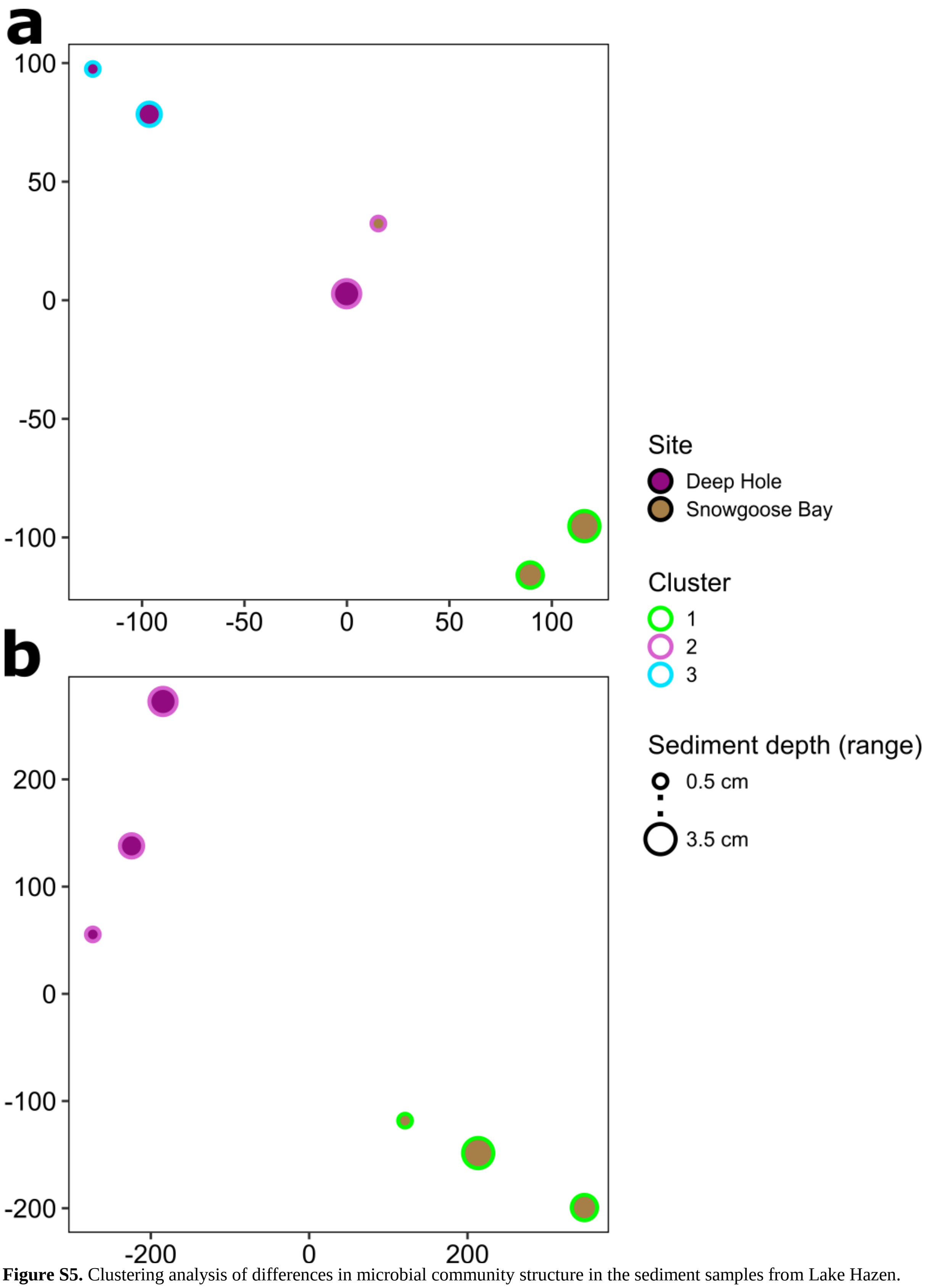

**Figure S5.** Clustering analysis of differences in microbial community structure in the sediment samples from Lake Hazen. (**a**) Medium quality MAGs ($n$ = 55) with manual taxonomy assignments based on reference genomes in a phylogenetic tree constructed from ribosomal proteins. (**b**) Raw reads binned into 16S rRNA contigs ($n$ = 166) with automatic taxonomy assignments from the SILVA 128 NR95 database.

#### 1.6 Marker genes and pathways

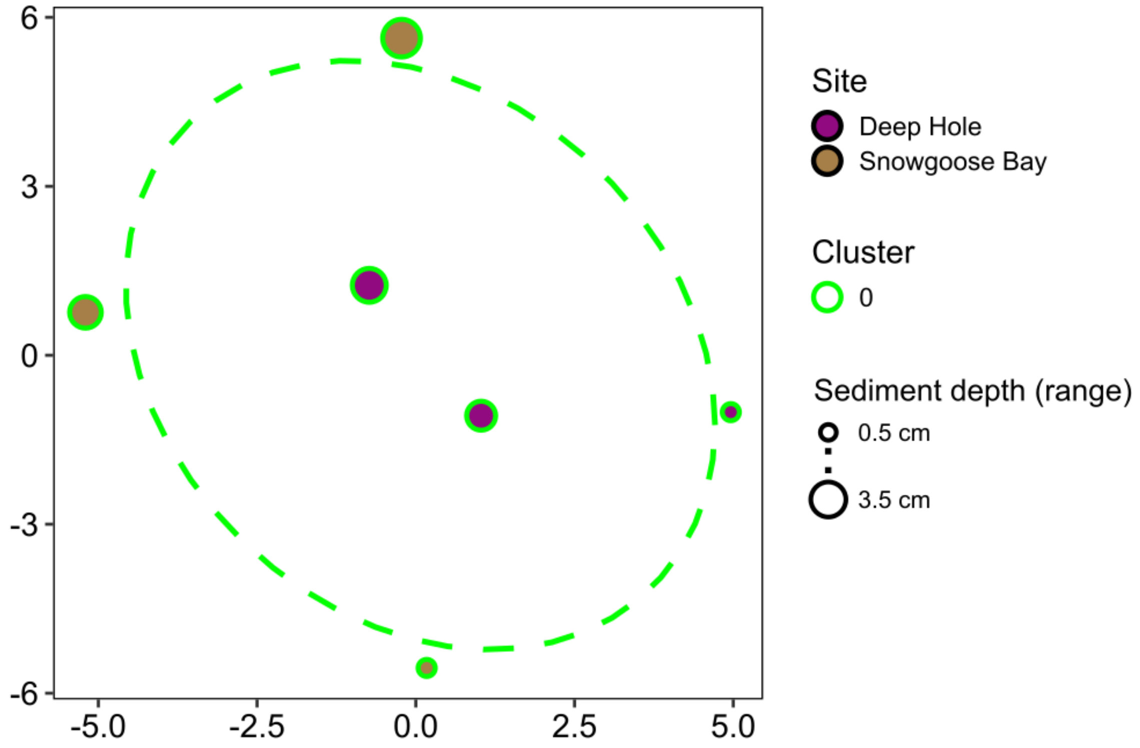

**Figure S6.** Clustering analysis of differences in MetaCyc functional pathway abundances in the 55 MAGs from sediment samples from Lake Hazen. All samples were assigned to 'Cluster 0' meaning that they are outliers, and no clusters were found in the analysis.

#### 1.7 Recently discovered and unfamiliar bacteria are found in Lake Hazen sediments

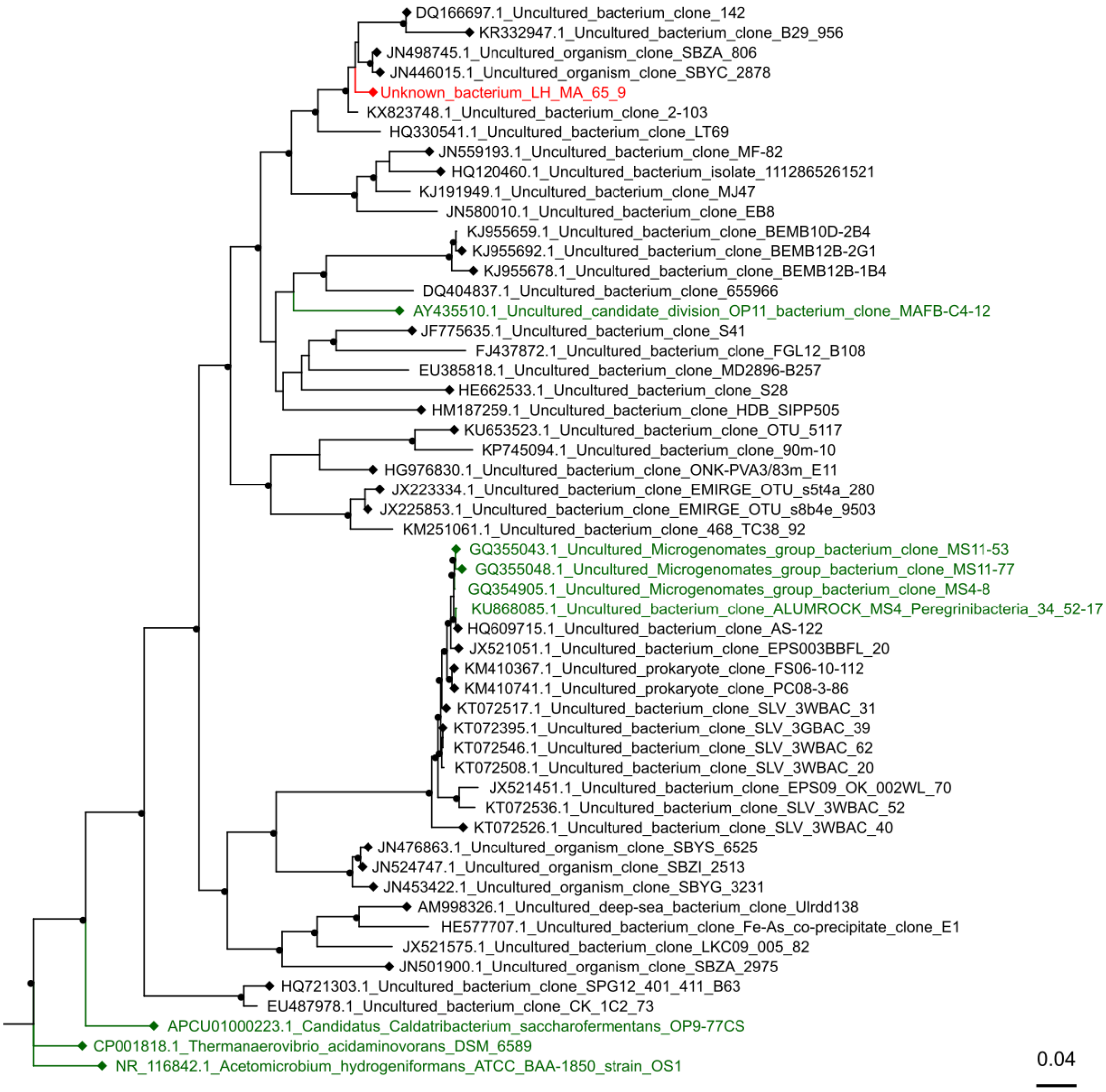

**Figure S7.** Phylogenetic tree of the taxonomically uncharacterized MAG, LH_MA_65_9, based on the alignment of its 16S rRNA gene. The MAG is shown in red and taxonomically classified 16S sequences in green. Circles on nodes indicate > 0.85 support in FastTree and diamonds on tips indicate higher than 90% nucleotide identity with the LH_MA_65_9 16S rRNA sequence over the aligned region. The tree was rooted with the 16S rRNA gene of *Acetomicrobium hydrogeniformans* (NCBI Reference Sequence: NR_116842.1).

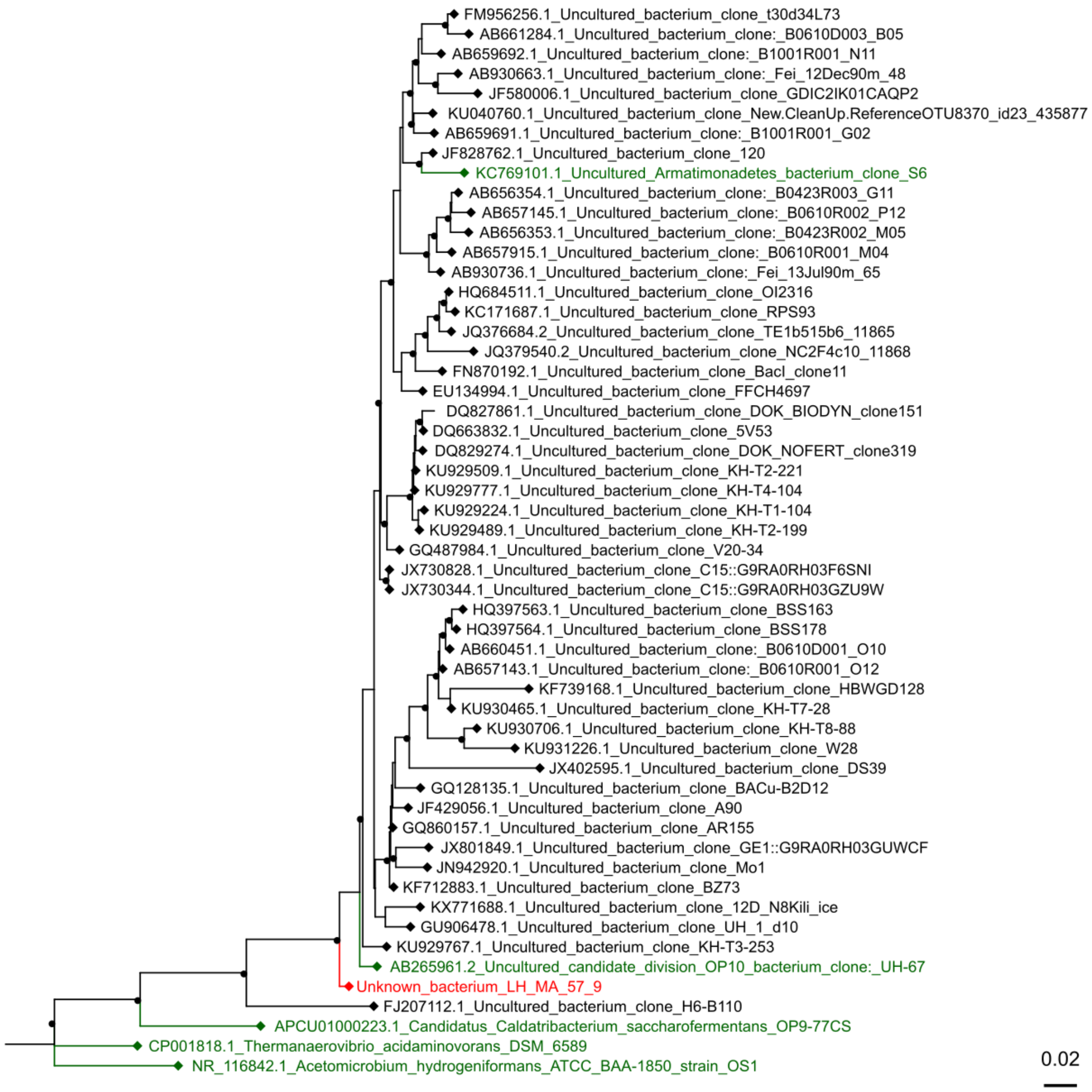

**Figure S8.** Phylogenetic tree of the taxonomically uncharacterized MAG, LH_MA_57_9, based on the alignment of its 16S rRNA gene. The MAG is shown in red and taxonomically classified 16S sequences in green. Circles on nodes indicate > 0.85 support in FastTree and diamonds on tips indicate higher than 90% nucleotide identity with the LH_MA_57_9 16S rRNA sequence over the aligned region. The tree was rooted with the 16S rRNA gene of *Acetomicrobium hydrogeniformans* (NCBI Reference Sequence: NR_116842.1).

#### 1.8 Nitrogen and sulfur cycles in the sediments

A Spirochaetes MAG was the most common nitrogen ($N_2$) fixer, together with three less abundant *Geobacter* MAGs (Figure 5a). Three Planctomycetes MAGs were the most common that harbored *nirAB* genes involved in Dissimilatory Nitrite Reduction to Ammonia (DNRA, $NO_2^-$ to $NH_4^+$). Planctomycetes MAGs were also the most common that had ammonia ($NH_4^+$) assimilation genes. Together, these three processes channel inorganic nitrogen into organic matter. Conversely, nitrogen is lost from aquatic systems by nitrification followed by denitrification, which releases it as NO, $N_2O$ or $N_2$ (Figure 5). Betaproteobacteria, specifically two Nitrosomonadales MAGs, were the only one which had the *haoB* gene involved in nitrification ($NH_4^+$ oxidation to $NO_2^-$), and the most abundant MAG harboring the *nosZ* marker gene for $N_2O$ reduction (to $N_2$), respectively. A *Rhodoferax* MAG (from Betaproteobacteria) was the most abundant nitrate ($NO_3^-$) reducer, harboring *nar*-genes. The ratio of DNRA to denitrification is important for the fate of nitrogen in the sediment as the former cycles nitrite back to ammonia and the latter reduces it to a volatile form. In Lake Hazen, 15 MAGs (27%) had genes involved in denitrification and 8 (15%) in DNRA. The mean relative abundances of the MAGs that had denitrification genes were also higher. However, the activity of these organisms in the two processes cannot be deduced purely from the analysis of metagenomic data, and marker genes for both processes were sometimes found in the same MAGs. A Myxococcales MAG (from Deltaproteobacteria) was the most abundant MAG with the *nrfH*-gene (involved in DNRA), but the organism was perhaps also capable of (*nirK*-mediated) nitrite reduction to the volatile nitrous oxide ($N_2O$), a key denitrification process. The relative contribution of these processes both in the community and in the same organism varies depending on the environment (van den Berg et al., 2017b) and can even be influenced by bacteria not directly involved in them (van den Berg et al., 2017a). It is not certain if denitrification (leading to losses of N) is more common in the Lake Hazen sediments. The lake appears to be a net sink of $NO_3^-$, $NO_2^-$ and $NH_4^+$ (St. Pierre et al., 2019), but because the top 1 to 2 cm of the sediment are oxidized (Figure S3), denitrification is unlikely to take place there. On the other hand, because of the high sediment deposition in Lake Hazen (St. Pierre et al., 2019), the nitrogen species that are deposited together with the particulate matter might get buried quite rapidly into the anoxic layers, where denitrification by these organisms could take place. However, further study would be required to assess the fate of nitrogen and its seasonal trends in Lake Hazen.

In the sulfur cycle, sulfate ($SO_4^{2-}$) in the sediment can either be reduced anaerobically by dissimilatory sulfate reducers to sulfite ($SO_3^{2-}$) or aerobically by assimilation into organic sulfur (Figure 5B). Marker genes for the dissimilatory pathway (*aprAB*) were detected in the > 1kb contigs, and the assimilatory pathway was detected in 10 of the 55 MAGs, with Planctomycetes as the most abundant phylum. The assimilatory pathway is likely utilized in the aerobic top sediments to synthesize cysteine and methionine, while the strictly anaerobic dissimilatory pathway is likely an electron sink deeper in the sediment. Sulfite can then be produced in the sediment directly by the dissimilatory pathway or desulfonated from organic sulfonates (Cook et al., 1998). Here, Alphaproteobacteria was the most abundant phylum with the desulfonation pathway. The 'NrfD-type' family (Pfam 03916) that we annotated as 'DNRA Polysulfide reductase' matches enzymes associated with both nitrite reduction to ammonia (Simon, 2002) and sulfur reduction (Jormakka et al., 2008). In Figure 5, we annotated polysulfide reduction with this gene rather than DNRA since we had more specific models for DNRA marker genes (namely *nrfH*). The marker gene for this more generic sulfur reduction (including tetrathionate-, DMSO-, and polysulfide-reductases) was found in several MAGs, of which Verrucomicrobia was the most abundant, followed by Deltaproteobacteria and Betaproteobacteria. The more specific *cysI*-mediated assimilatory sulfite reduction marker was only found in two Verrucomicrobia bacteria (also harboring the other sulfur reduction marker gene).

Only few MAGs had markers for multiple sulfur reduction pathways, and none could probably catalyze the complete reduction of sulfate to sulfide. Furthermore, we could not find the markers for *dsrAB* or *aprAB* genes from the MAGs while they were found, respectively, from 61 and 54 repository genomes using the same pipeline. They were also found in the low-quality bins and unbinned contig data (Figure 4). The *dsrAB* genes encode proteins that catalyze the reduction of sulfite to sulfide, or act in reverse in the oxidation of sulfide (Müller et al., 2015). The *aprAB* genes similarly encode for proteins catalyzing the reversible reduction of sulfate to sulfite (Rabus et al., 2006). The absence of these pathways in the 55 MAGs shows that these pathways are likely rare, but definitely not absent in the Lake Hazen sediments. Finally, thiosulfate is also oxidized in the sediment; a single MAG from Nitrosomonadales contained the *soxAXYZ*-genes.

#### 1.9 Nutrient cycling capabilities of individual MAGs

Certain MAGs appeared to have highly versatile metabolisms. The only MAG with marker genes for thiosulfate oxidation (*soxAXYZ*) from the order Nitrosomonadales (LH_MA_55_1) likely represents an organism capable of assimilatory sulfite reduction to sulfide (as it has a *cysI* gene) and is also important in nitrogen cycling (Table S6): the MAG contained both a respiratory nitrate reductase (*narI*) and a nitrite reductase (*nirBD*), which produce ammonium ($NH_4^+$) as the product. The MAG also contained a nitrous oxide reductase (*nosZ*), which could be used for respiration. Furthermore, the MAG of this likely autotrophic organism contained a wide array of biosynthetic pathways, e.g., a pentose phosphate pathway, and a reverse TCA cycle likely utilized for carbon fixation. A *Geobacter* (LH_MA_37_3) MAG included marker genes for N fixation (*nifS*, *nifV*), ammonia assimilation (*gltB*), nitrate reduction to nitrite (*narGHIJ*), DNRA / Polysulfide reduction (*nrfD*), and likely has wide biosynthetic capabilities. Similarly, two Chloroflexi (LH_MA_58_17 and LH_MA_61_3) MAGs and a Spirochaete (LH_MA_58_12) MAG had several marker genes for both nitrogen and sulfur cycle processes. The MAG LH_MA_65_9, which was unclassified at the phylum-level, can likely both fix nitrogen (*nifS*) and utilize nitrite through DNRA (*nrfH*) and ammonia assimilation (*amtB*, *gltB*). None of the CPR MAGs had any of the nutrient cycling marker genes or MetaCyc pathways.

### References

Cook, A. M., Laue, H., and Junker, F. (1998). Microbial desulfonation. *FEMS Microbiol. Rev.* 22, 399–419. doi:10.1111/j.1574-6976.1998.tb00378.x.

Hurt, R. A., Qiu, X., Wu, L., Roh, Y., Palumbo, A. V., Tiedje, J. M., et al. (2001). Simultaneous recovery of RNA and DNA from soils and sediments. *Appl. Environ. Microbiol.* 67, 4495–4503. doi:10.1128/AEM.67.10.4495-4503.2001.

Jormakka, M., Yokoyama, K., Yano, T., Tamakoshi, M., Akimoto, S., Shimamura, T., et al. (2008). Molecular mechanism of energy conservation in polysulfide respiration. *Nat. Struct. Mol. Biol.* 15, 730–737. doi:10.1038/nsmb.1434.

Köck, G., Muir, D., Yang, F., Wang, X., Talbot, C., Gantner, N., et al. (2012). Bathymetry and sediment geochemistry of Lake Hazen (Quttinirpaaq National Park, Ellesmere Island, Nunavut). *Arctic*, 56–66.

Müller, A. L., Kjeldsen, K. U., Rattei, T., Pester, M., and Loy, A. (2015). Phylogenetic and environmental diversity of DsrAB-type dissimilatory (bi)sulfite reductases. *ISME J.* 9, 1152–1165. doi:10.1038/ismej.2014.208.

Rabus, R., Hansen, T. A., and Widdel, F. (2006). Dissimilatory sulfate- and sulfur-reducing Prokaryotes. *The Prokaryotes*, 659–768. doi:10.1007/0-387-30742-7_22.

Simon, J. (2002). Enzymology and bioenergetics of respiratory nitrite ammonification. *FEMS Microbiol. Rev.* 26, 285–309. doi:10.1111/j.1574-6976.2002.tb00616.x.

St. Pierre, K. A., St. Louis, V. L., Lehnherr, I., Schiff, S. L., Muir, D. C. G., Poulain, A. J., et al. (2019). Contemporary limnology of the rapidly changing glacierized watershed of the world's largest High Arctic lake. *Sci. Rep.* doi:10.1038/s41598-019-39918-4.

van den Berg, E. M., Elisário, M. P., Kuenen, J. G., Kleerebezem, R., and van Loosdrecht, M. C. M. (2017a). Fermentative bacteria influence the competition between denitrifiers and DNRA bacteria. *Front. Microbiol.* 8. doi:10.3389/fmicb.2017.01684.

van den Berg, E. M., Rombouts, J. L., Kuenen, J. G., Kleerebezem, R., and van Loosdrecht, M. C. M. (2017b). Role of nitrite in the competition between denitrification and DNRA in a chemostat enrichment culture. *AMB Express* 7. doi:10.1186/s13568-017-0398-x.
