## Supplementary figures and images for "Microbial genomes retrieved from High Arctic lake sediments encode for adaptation to cold and oligotrophic environments"

### Full_tree_with_supports.pdf

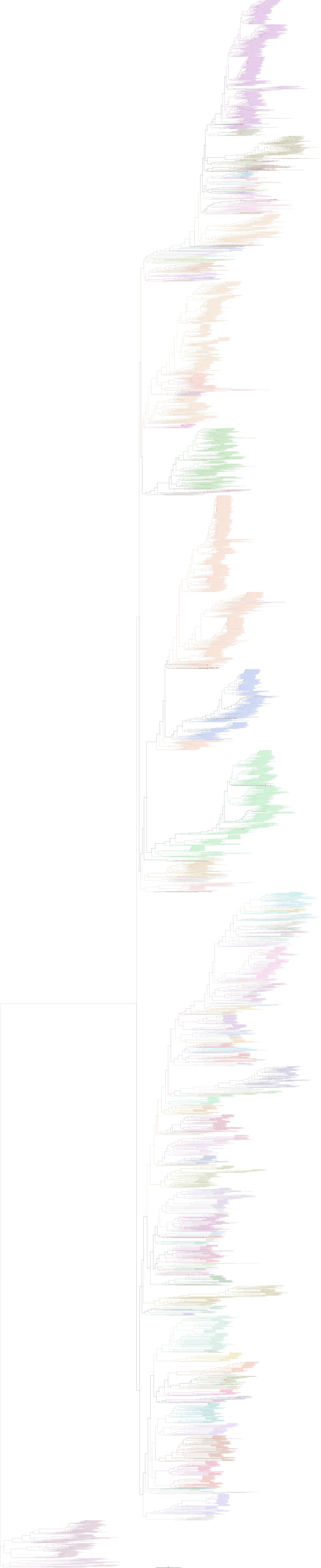
